# Forecasting of phenotypic and genetic outcomes of experimental evolution in *Pseudomonas protegens*

**DOI:** 10.1101/342261

**Authors:** Jennifer T. Pentz, Peter A. Lind

## Abstract

Experimental evolution with microbes is often highly repeatable under identical conditions, suggesting the possibility to predict short-term evolution. However, it is not clear to what degree evolutionary forecasts can be extended to related species in non-identical environments, which would allow testing of general predictive models and fundamental biological assumptions. To develop an extended model system for evolutionary forecasting, we used previous data and models of the genotype-to-phenotype map from the wrinkly spreader system in *Pseudomonas fluorescens* SBW25 to make predictions of evolutionary outcomes on different biological levels for *Pseudomonas protegens* Pf-5. In addition to sequence divergence (78% amino acid and 81% nucleotide identity) for the genes targeted by mutations, these species also differ in the inability of Pf-5 to make cellulose, which is the main structural basis for the adaptive phenotype in SBW25. The experimental conditions were also changed compared to the SBW25 system to test the robustness of forecasts to environmental variation. Forty-three mutants with increased ability to colonize the air-liquid interface were isolated, and the majority had reduced motility and was partly dependent on the *pel* exopolysaccharide as a structural component. Most (38/43) mutations are expected to disrupt negative regulation of the same three diguanylate cyclases as in SBW25, with a smaller number of mutations in promoter regions, including that of an uncharacterized polysaccharide operon. A mathematical model developed for SBW25 predicted the order of the three main pathways and the genes targeted by mutations, but differences in fitness between mutants and mutational biases also appear to influence outcomes. Mutated regions in proteins could be predicted in most cases (16/22), but parallelism at the nucleotide level was low and mutational hot spots were not conserved. This study demonstrates the potential of short-term evolutionary forecasting in experimental populations and provides testable predictions for evolutionary outcomes in other *Pseudomonas* species.

**Author Summary:** Biological evolution is often repeatable in the short-term suggesting the possibility of forecasting and controlling evolutionary outcomes. In addition to its fundamental importance for biology, evolutionary processes are at the core of several major societal problems, including infectious diseases, cancer and adaptation to climate change. Experimental evolution allows study of evolutionary processes in real time and seems like an ideal way to test the predictability of evolution and our ability to make forecasts. However, lack of model systems where forecasts can be extended to other species evolving under different conditions has prevented studies that first predict evolutionary outcomes followed by direct testing. We showed that a well-characterized bacterial experimental evolution system, based on biofilm formation by *Pseudomonas fluorescens* at the surface of static growth tubes, can be extended to the related species *Pseudomonas protegens*. We tested evolutionary forecasts experimentally and showed that mutations mainly appear in the predicted genes resulting in similar phenotypes. We also identified factors that we cannot yet predict, such as variation in mutation rates and differences in fitness. Finally, we make forecasts for other *Pseudomonas* species to be tested in future experiments.

## Introduction

An increasing number of experimental evolution studies, primarily using microbes, have provided insights into many fundamental questions in evolutionary biology including the repeatability of evolutionary processes (1-6). Given the ability to control environmental conditions, population size, as well as the use of a single asexual organism, such studies could provide an ideal test of our ability to predict evolutionary outcomes in simplified model systems. High repeatability on both phenotypic and genetic levels have been observed in a large number of experimental evolution studies (reviewed in (5)), but it has become clear that high repeatability alone is not sufficient for testing evolutionary predictability beyond the prediction that under identical conditions the same evolutionary outcome is probable.

The difficulties of moving from repeatability to predictability are largely a result of the lack of knowledge of the genotype-phenotype-fitness map, including how sensitive it is to changes in environmental conditions and to what degree it is conserved between different strains and species (7-9). Several problems arise when searching for suitable model systems for testing and improving predictive ability (reviewed in detail in (10)): I. Adaptive mutations are often highly strain specific, so that adaptation of different strains of the same species to a specific environment will produce different results (6, 11). II. The range of probable adaptive phenotypes can often not be defined beforehand due to more than one dominant selective pressures (6, 12, 13), which means that in many cases the phenotypes that solve the focal selective problem are outcompeted by other phenotypes with increased fitness. III. Adaptation to a specific selective pressure may be solely explained by the molecular phenotype of a single protein, resulting in a relatively simple parameter and genotype space, as is sometimes observed for high-level antibiotic resistance (14-19). Thus, predictions will be identical for all species and the model system cannot provide a test of predictions from general principles. The most useful model systems for testing and developing our ability to predict evolution, at least at this point, are likely to be of intermediate complexity with several beforehand recognizable phenotypic and genetic solutions to combine ample opportunities for failure with a decent chance of success.

The wrinkly spreader (WS) model in *P. fluorescens* SBW25 (hereafter SBW25) is one of the most well characterized experimental evolution systems and has several suitable properties, in relation to the problems described above, that could make it possible to extend knowledge and principles from this species to related species (20-27). When the wild type SBW25 is placed into a static growth tube the oxygen in the medium is rapidly consumed by growing bacteria. However, the oxygen level at the surface is high and mutants that can colonize the air-liquid interface have a major growth advantage and rapidly increase in frequency (Fig 1A). Several phenotypic solutions to air-liquid interface colonization, all involving increased cell-cell adhesion, have been described and are distinguishable by their colony morphology on agar plates (20, 22, 25). The most successful of these is the Wrinkly Spreader (WS) (20, 22) that overproduces cellulose that is the main structural component of the mat at the air-liquid interface (24, 28). The WS phenotype is caused by mutational activation of c-di-GMP production by a diguanylate cyclase (DGC) (27). While many different DGCs can be activated to reach the WS phenotype, some are greatly overrepresented due to larger mutational target sizes leading to a hierarchy of genetic routes to WS (Fig 1D) (21, 23). The genotype-to-phenotype map to WS has been characterized in detail (23, 26, 27) allowing the development of mathematical models of the three main pathways to WS (Wsp, Aws and Mws) and the prediction of evolutionary outcomes (26).

**Fig. 1.**
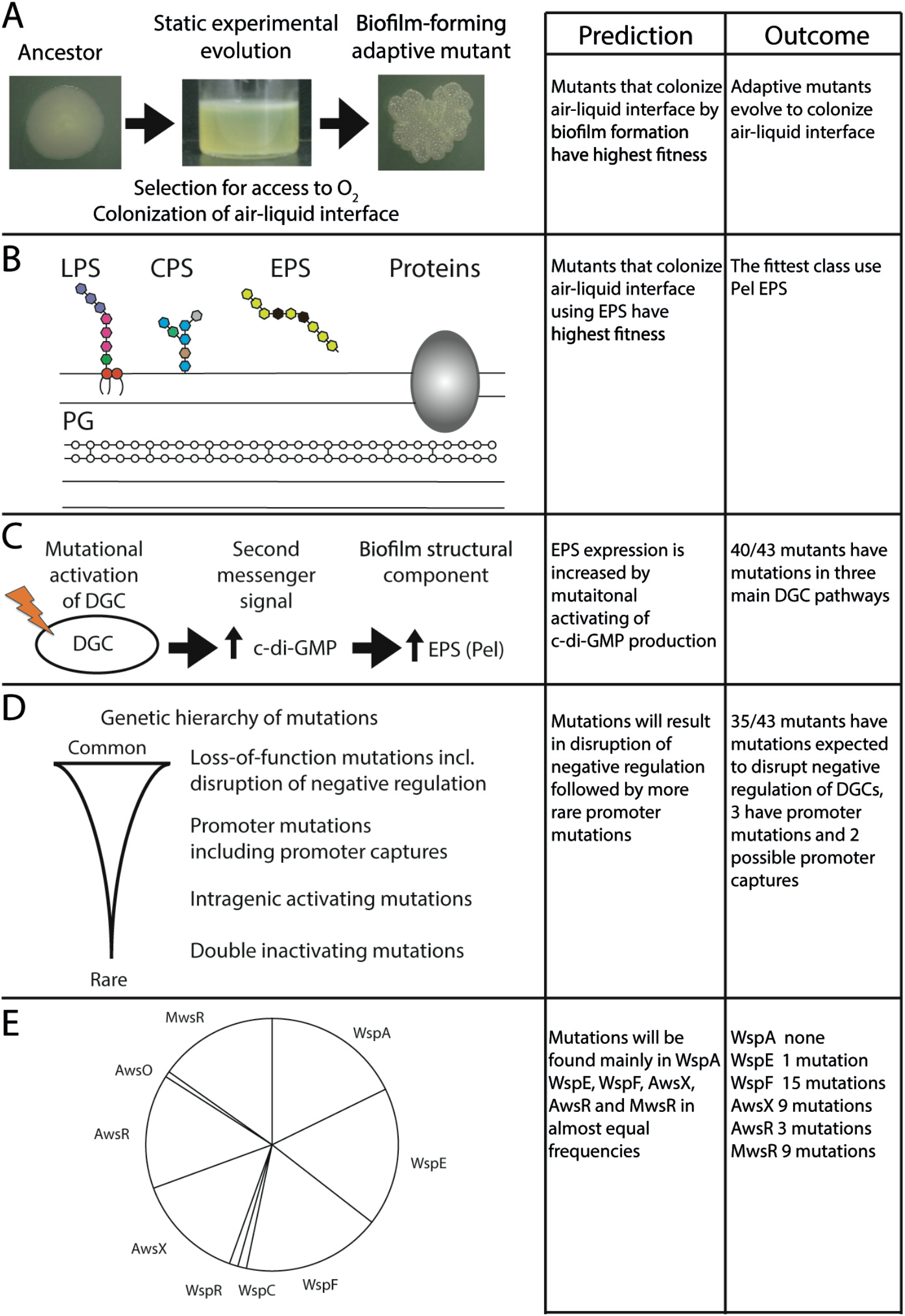
Evolutionary predictions and outcomes of experimental evolution for *Pseudomonas protegens* Pf-5. (*A*) Growth under static conditions when oxygen is limiting to growth is expected to result in selection for colonization of the air-liquid interface by increased cell-cell adhesion and surface attachment. Pictured is a wrinkly spreader mutant of *Pseudomonas fluorescens* SBW25. (*B*) The cell wall of Pf-5 has several components that could possibly be used to promote cell-cell adhesion and surface attachment, including lipopolysaccharides (LPS), capsular polysaccharides (CPSs), exopolysaccharides (EPSs), adhesive proteins, and incomplete cleavage of the peptidoglycan (PG) layer. (*C*) Overexpression of an EPS caused by mutational activation of a di-guanylate cyclase (DGC) leading to increased c-di-GMP production is predicted to produce the fittest phenotypes. Furthermore, increased c-di-GMP is also predicted to result in reduced motility. (*D*) Based on previous experiments in SBW25 (21), the types of adaptive mutations expected for Pf-5 include, in decreasing frequency, loss-of-function mutations, promoter mutations, intragenic activating mutations, and double inactivating mutations. (*E*) Predicted fraction of WS mutations for genes in the main three pathways. Knowledge of the molecular networks allowed for the formulation of a mathematical model that accurately predicted the relative rates of use of the common three pathways (Wsp, Aws, and Mws) (26) and rates for genes in each pathway. The assumption that the molecular functions of these networks are conserved in Pf-5 allows prediction of the rates of pathways and genes.

In this study, we use previous knowledge of the SBW25 WS system to make predictions of phenotypic and genetic evolutionary outcomes of experimental evolution for *Pseudomonas protegens* Pf-5 (hereafter Pf-5) that are then tested experimentally. Pf-5 has a highly conserved genetic repertoire of DGCs, but lacks the main structural component used for air-liquid interface colonization by WS types in SBW25. Results show that phenotypes, order of pathways used, and types of mutations can be predicted and that forecasts are robust to introduced changes in experimental conditions compared with the SBW25 system. Our results suggest that there is potential for forecasting short-term evolutionary processes in an extended *Pseudomonas* wrinkly spreader system, and we conclude by making forecasts for five other species of *Pseudomonas* based mainly on their genome sequence and data from Pf-5 and SBW25 that will be tested in future studies (20-23, 29).

## Results

We aim to explore the potential of developing the wrinkly spreader system as a model system for testing evolutionary forecasts and developing methodology by first making predictions on several biological levels followed by experimental tests. Thus, the aim here is not to replicate wrinkly spreader experimental evolution in other species, but rather to see to what degree it fulfills the requirements of a suitable experimental system for testing evolutionary forecasts. One of these requirements is that a dominant selective pressure, access to oxygen at the air-liquid interface, can be established that is not dependent of the specific experimental conditions. Therefore we changed conditions compared to the SBW25 systems in terms of growth medium (KB changed to TSB with glycerol and MgSO_4_), temperature (28°C to 36°C), growth vessel (glass tube to polypropylene deep well plate) and final population size (about 2 × 10^10^ to 4 × 10^8^). As a first test we chose the related species *P. protegens* Pf-5 for a parallel experimental evolution study under static conditions (Fig 1A). This strain has all the main genetic pathways to WS conserved, with an average of 78% amino acid and 81% nucleotide identity (S1 Table) for the genes commonly targeted by mutations, while lacking the genes for cellulose biosynthesis, which is the main structural component used for colonization of the air-liquid interface in SBW25. This makes Pf-5 suitable for an initial test of genetic robustness, *i*.*e*. that species with similar genetic composition adapt using the same genetic pathways. No previous data on WS mutations or phenotypes with the ability to colonize the air-liquid interface have been published for P. *protegens* Pf-5 to our knowledge.

### Predictions for experimental evolution of *P. protegens* Pf-5

All evolutionary forecasts are dependent on previous knowledge to some degree, in terms of theory and empirical data, combined with an assessment of how likely these are to be applicable to the specific novel situation. Therefore it is necessary to describe the bases for predictions on different biological levels together with possible alternative outcomes to be able to judge the underlying assumptions and potential biases influencing the forecasts and to determine why predictions fail.

Forecasting experimental evolution in a novel context (*e*.*g*., a new species or different environment) first requires a prediction of the dominant selective pressures and the corresponding phenotypes that will have the highest fitness. Only then can more specific phenotypic predictions be made, followed by genetic predictions when the genotype-to-phenotype map is well characterized. Thus, predictions of specific phenotypes are dependent on a correct prediction of dominant selective pressures and any genetic predictions will be dependent on correct phenotypic predictions. Therefore, we first describe predictions on different biological levels in the order of specialization, phenotypes, genetics and molecular details. However, analysis of the outcomes of experimental evolution will often start with identifying the causal mutations followed by reconstruction into the ancestral genetic background to determine the effects of mutations on the phenotypic level. Therefore, conversely to describing the evolutionary predictions, experimental results are presented starting with mutational data, followed by characterization of phenotypes and fitness. For a quick reference, a summary of predictions and outcomes on different biological levels is available in Fig 1.

### Mutants that specialize in colonizing the air-liquid interface will have the highest fitness

It is likely that there are several evolutionary strategies with increased fitness under the experimental conditions used here, but we predict that selection for access to oxygen will be the dominant selective pressure. Mutants with an increased ability to colonize the air-liquid interface will be the fittest class of adaptive mutants and rapidly increase in frequency due to the large increase in growth rate under high oxygen conditions (Fig 1A). This is likely to be true only when the wild type is a poor colonizer of the air-liquid interface. The cell wall of Gram-negative bacteria, such as *Pseudomonas*, has several components that may be used to increase cell-to-cell adhesion and attachment to surfaces to allow colonization of the air-liquid interface (Fig 1B). Thus, all bacteria that are obligate aerobes, like Pf-5, are predicted to be able to mutate to find a phenotypic solution to colonize the air-liquid interface under conditions where selection for access to oxygen is strong.

### Non-motile mutants with increased production of exopolysaccharides will form the most stable mats at the air-liquid interface and outcompete other phenotypic solutions

If mutants specializing in colonizing the air-liquid interface are the fittest class, we can make a prediction of the specific phenotypic solutions used for increasing cell-to-cell and/or cell-surface adhesion. A key parameter here is the stability of the mat, as it will collapse and fall to the bottom when a self-supporting mat covering the surface cannot be maintained. In addition to different exopolysaccharides (EPSs), capsular polysaccharides (CPSs) and lipopolysaccharides (LPS) might also be used (Fig 1B). Adhesive proteins and incomplete cleavage of the peptidoglycan (PG) layer are also potential phenotypic solutions for increased cell-cell adhesion. In SBW25, at least four alternative distinct phenotypes are selected for at the surface, involving an alternative EPS, CPS, LPS and PG, but they all have lower fitness than the WS type that form cellulose-based mats (20, 22, 30, 31). Pf-5 lacks genes for cellulose biosynthesis, but there are other EPSs that might be used (*e*.*g*., Pel, Psl, polyglucosamine (PGA)). Thus, we predict that overexpression of an EPS will always provide the most stable mats and therefore have the highest fitness (Fig 1C) and that Pel will be the primary EPS used based on its importance for pellicle formation at the air-liquid interface in *Pseudomonas aeruginosa* PA14 (32). This prediction might fail, though, because the encoded EPS does not form superior mats, is non-functional, or the biosynthetic costs of EPS production outweighs its benefit relative to other phenotypic solutions.

In SBW25 and *Pseudomonas aeruginosa* PA14, overexpression of EPSs used for mat formation are linked to mutations increasing c-di-GMP production rather than mutations in the promoters of, or genes in, the EPS operons themselves (21, 22, 33). This can be explained by the role of post-translation regulation by c-di-GMP in the production of the EPSs cellulose, Pel, PGA and alginate (34). Furthermore, there may be an additional benefit of using c-di-GMP activation in that it reduces motility, which is not needed when bacteria are established at the air-liquid interface and may be antagonistic to biofilm formation (35). Motility also consumes a large amount of energy and thus is likely to be lost when not under selection (36, 37). Consequently, we also predict that the fittest mutants overexpressing EPSs will have reduced motility (Fig 1C).

### Genetic predictions

Predictions on the genetic level are more difficult depending on the greater dimensionality of the problem (*i*.*e*., mutations in many genes can lead to the same phenotype) but also because genetic predictions will be conditional on a correct phenotypic prediction. The genotype-to-phenotype map must also be conserved to some degree between species, so that similar mutations give rise to similar, previously observed, phenotypes. Mutational biases can also greatly skew the diversity of mutants found for a particular species, which can cause predictions to fail even if the theoretical model of the genotype-to-phenotype map is accurate. However, the ability to construct predicted mutants that are not found after experimental evolution provides the possibility to test if the mutation produces the expected phenotype and measure its fitness against the mutants found in experimental evolution. This provides an opportunity to determine the underlying causes of why certain predicted mutants are not observed after experimental evolution, which can lead to improved predictive models.

In cases where the phenotypic prediction fails, for example because a novel previously unknown phenotypic solution to the adaptive problem is used, or the genotype-to-phenotype map is not conserved, detailed genetic predictions will inevitably fail. However, a general prediction of the types of mutations is still possible based on empirical observations. Loss-of-function mutations account for a majority of adaptive mutations in many experimental evolution studies (38), presumably because of a large mutational target size. This is true also for SBW25 where a large majority of WS mutations are loss-of-function mutations in negative regulators (WspF, AwsX) or intragenic negative regulatory regions (MwsR, PFLU0085) of the main DGCs (23) as well as loss-of-function mutations in the genes underpinning the alternative phenotypes (20, 22, 23). When the negatively regulated pathways to WS are removed, DGCs are instead activated by promoter mutations, including promoter captures followed by more rare intragenic activating mutations (21) (Fig 1D).

#### Prediction of DGC pathways and mutated genes by modeling of the WS phenotype-to-genotype map

If WS types in Pf-5 evolve using mutational activation of DGCs, as expected if the genotype-to-phenotype map is conserved, we can use previously developed mathematical models (26) to predict the relative rates for the three main pathways (Wsp, Aws, and Mws) to the WS phenotype based on a detailed understanding of the molecular functions of the proteins involved (Fig 1E, S1 Text). The molecular networks of the three main pathways (Wsp, Aws, and Mws) were modeled as a system of ordinary differential equations (S1 Text, S1 Fig, S2 Fig) and this work is fully described as Model IV in Lind et al. 2019 (26). The functional interactions between components of the networks, for example an enzymatic reaction or a conformational change in a protein, are described by biochemical reaction rates that can be either increased or decreased by mutations, resulting in changes in the concentration of the active form of the DGC of each pathway. We assume here that the functions of the proteins in the Wsp, Aws, and Mws regulatory networks are conserved between SBW25 and Pf-5. We do not assume that the concentrations of the proteins or the biochemical reaction rates are identical as these are unknown for both species and are repeatedly drawn from uniform distributions. This allows us to estimate, by numerical simulations, the relative probability that mutational changes in reaction rates for each pathway result in a phenotypic change in the form of a wrinkly spreader. The model is independent of the exopolysaccharide used as the structural component, as it only calculates the probability that a mutation increases the active form of the DGC for each pathway. The model can also incorporate information about mutational biases by increasing probability of changes in specific reactions rates, which improved the predictive ability of the model in Lind et al. 2019. We do not assume that mutational hot spots are conserved and therefore all reaction rates had the same probability of mutational changes. Based on data from Lind et al. 2019 and the general expectation that disabling mutations (reducing reaction rates) are more likely than enabling mutations (increasing reaction rates), we assumed that disabling changes are ten times more common (Fig 1E).

The mathematical model of the main DGC pathways can also predict the relative contribution of changes to each reaction rate to phenotypic change ((26), S1 Text). With an understanding of the functions of the proteins involved this allows prediction of functional effects of mutations and thereby also targeted regions in the proteins. Here we make the simplifying assumption that changes to a reaction rate is equally likely to be caused by mutations in all participating proteins. With more data on mutational target size of each protein it would be possible to integrate this information, which might improve future predictions. High rates of WS mutations are predicted for WspF, WspA, WspE, AwsX and AwsR and MwsR (Fig 1E). A significantly lower rate of enabling mutations is also predicted to occur in WspC, WspR and AwsO (Fig 1E). Despite the simplicity of the null model, it closely predicted the mutational targets in SBW25 with equal rates for WspF, WspA and WspE and rare mutations in WspC and WspR, suggesting that it is a useful null model for other species (26).

#### Prediction of specific mutational targets and effects of mutations

If the genotype-to-phenotype map is conserved between SBW25 and Pf-5 and modeling allows prediction of mutated genes and their functional effects on reaction rates, we can proceed to make more detailed predictions about the specific mutations for each gene. The level of parallelism at the nucleotide level between species is expected to be dependent on the number of possible mutations to WS and the degree of functional conservation of the proteins involved that define the genotype-to-phenotype map. For Pf-5, the genes in the four main pathways (Wsp, Aws, Mws, PFLU0085/DgcH) have a nucleotide identity of 71-84% and an amino acid identity of 68-92% (S1 Table). Thus, identical base pair substitutions are not always possible and, without studying each single case, the probability of parallelism at the nucleotide or amino acid level cannot be calculated and is not included in our predictions. Mutational hot spots with greatly increased mechanistic mutation rates are also expected to contribute to parallelism when they are conserved, but reduce parallelism when they are not. Based on previous analysis of patterns of mutations in SBW25 (23, 26, 29) and homology modeling of protein structure using Phyre2 (39), regions expected to be mutated were predicted and the likely molecular consequences of different mutations suggested (Fig 2). The mutational effects on protein function can be directly connected to the mathematical model of the genotype-to-phenotype map by linking their likely effects to increasing or decreasing reaction rates (S1 Text).

**Fig. 2.**
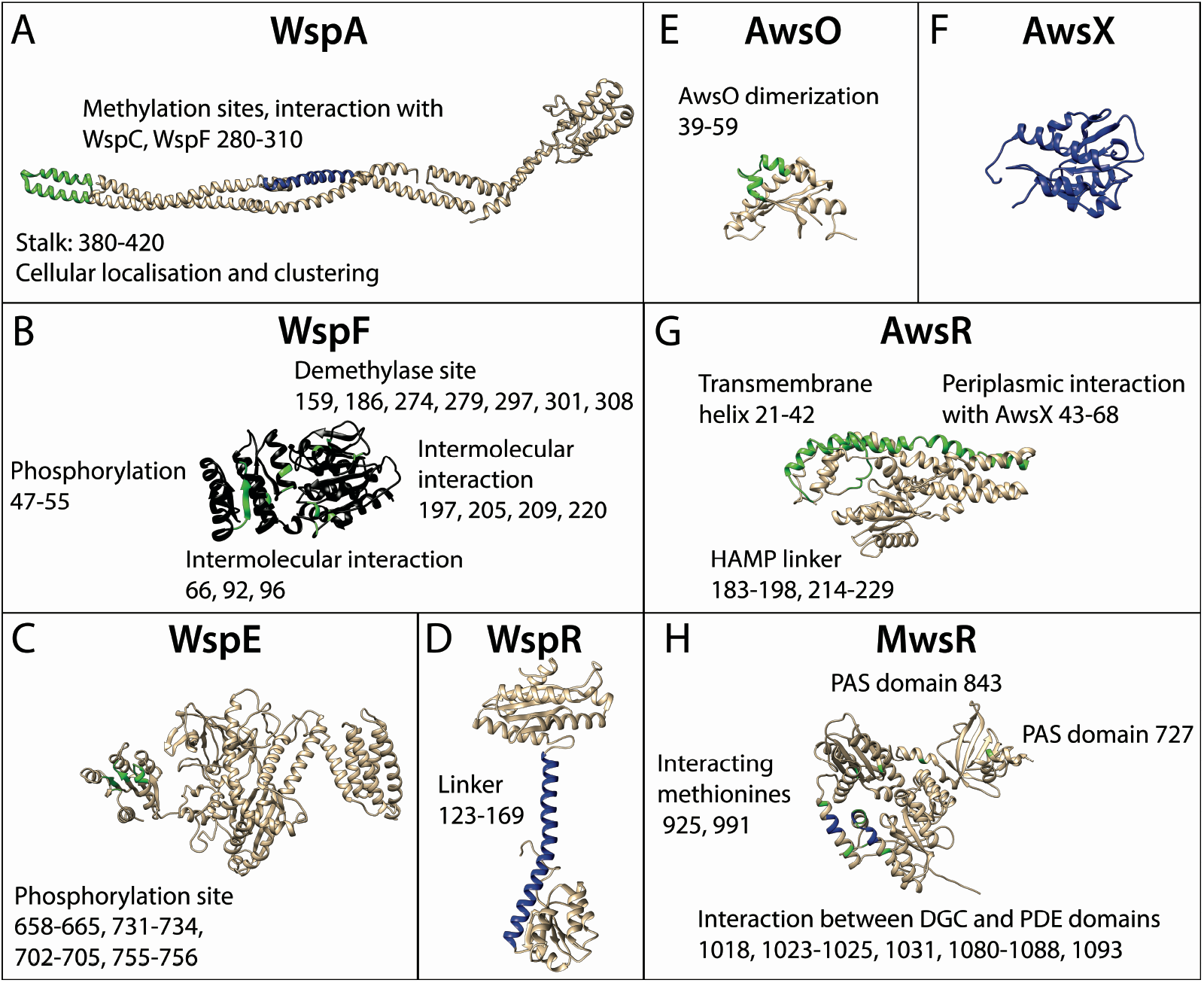
Predicted mutational targets and proposed molecular effects. Black represents any inactivating mutation including frame shifts, blue represents in frame inactivating mutations, green represents amino acid substitutions. Numbers refer to amino acid residue numbers in Pf-5. (*A*) WspA – amino acid substitutions are expected at the tip of the stalk and in-frame deletion of methylation sites. (*B*) WspF – any inactivating mutation is predicted, amino acid substitutions are predicted only in areas where they disrupt intermolecular interactions. (*C*) WspE – amino acid substitutions are predicted near the phosphorylation site. (*D*) WspR – small in frame deletion and amino acid substitutions in the linker is predicted to cause constitutive activation. (*E*) AwsO – amino acid substitutions disrupting AwsO dimerization is predicted to lead to increased binding to AwsX without the presence of an activating signal. (*F*) AwsX – any inactivating mutation that keep the reading frame intact and do not interfere with expression of downstream AwsR is predicted. (*G*) AwsR - amino acid substitutions in the periplasmic region or transmembrane helix that disrupt the interaction with AwsX or to the HAMP linker is predicted. (*H*) MwsR – mutations are predicted in the interface between the DGC and phosphodiesterase domains and in the most C-terminal of the PAS domains resulting in constitutive activation.

### Prediction of fitness effects of WS mutations

Even if the model of the genotype-to-phenotype map can accurately predict the rates of different WS mutants for Pf-5, previous results show that some WS mutants have lower fitness and are rarely observed after experimental evolution (21, 26). Thus, in order to produce more accurate predictions there is a need to *a priori* predict the relative fitness of different WS mutants using previous experimental data, assuming these will be conserved between species, or by theoretical considerations. The distribution of fitness effects of advantageous mutations has mainly been explored theoretically by Gillespie (40-42) and Orr (43, 44), but this work assumes that the wild type is well adapted, which is not the case here as the wild type is a poor colonizer of the air-liquid interface. Noting that the most common mutations observed in SBW25 are likely to disable intermolecular or interdomain interactions (26) we instead turn to the literature on the distribution of fitness effects of mutations that *reduce* function and are therefore typically deleterious. The distribution of fitness effects of random mutations have been found to be bimodal for a large number of genes with different functions, with one mode close to neutrality and one corresponding to a complete loss of a particular molecular function (45-49). In cases where disabling mutations in a gene are advantageous, we expect a similar bimodal distribution but that the second mode is centered at higher fitness than the wild type. However, there is no reason to expect that the mode of disabling mutations in different genes to be centered at the same fitness, which means that the observed WS mutants will be biased towards the genes with the highest fitness modes rather than distributed equally among them. The overall conclusion is that even if there are many possible genetic pathways to an adaptive phenotype in experimental evolution, we predict that mutations will only be found in the subset of genes with modes at highest fitness. If the ranks of relative fitness of the modes are conserved between species this would greatly aid predictions, but if they are not limited experimental data will be needed to predict the high fitness genes.

### Experimental test of forecasts in *Pseudomonas protegens* Pf-5

To determine if the WS system can be extended to a related species and to experimentally test evolutionary forecasts in Pf-5, we inoculated 60 independent wells with the wild type for five days of experimental evolution under static conditions. Air-liquid interface colonization was observed for the majority of the wells and independent mutants with clearly visible changes in colony morphology were observed for 43 wells. One colony for each well was selected for further characterization at random based on a pre-determined position on the agar plate.

#### The majority of mutations are found in the Wsp, Aws and Mws pathways

We started with evaluating our predictions on the genetic level, in terms of types of mutations, DGC pathways and mutational targets, as it is relatively straightforward to find mutations using DNA sequencing. Identifying the causal mutations also provides clues to the phenotypic basis of air-liquid interface colonization and is needed to be able to reconstruct mutations to determine causality and measure fitness in the absence of possible secondary mutations.

The causal mutations were identified using a combination of Sanger and Illumina genome sequencing (Fig 3, S2 Table). As predicted, the majority (40/43) of mutations were associated with the Wsp, Aws, and Mws DGC pathways (Fig 1C) that are subject to negative regulation (Fig 1D). In addition, the prediction that promoter mutations would be the second most common type of mutation (Fig 1D) was accurate, with two mutations found upstream of the *aws* operon that were predicted to disrupt the terminator of a high expression ribosomal RNA operon representing a putative promoter capture event. Promoter mutations were also found upstream of PFL_3078, which is the first gene of a putative polysaccharide locus (PFL_3078-3093) that has recently been characterized *in silico* and named *Pseudomonas* acidic polysaccharide (Pap) (50). Pap is a newly discovered polysaccharide encoded by PSF113_1955-1970 in *Pseudomonas fluorescens* F113 (50), which is only present in closely related strains (and not in SBW25). The operon encodes genes typical and necessary for the biosynthesis of a polysaccharide (50) and it is likely that PFL_3078-3093 encodes the main biofilm structural component used by these mutants.

**Fig. 3.**
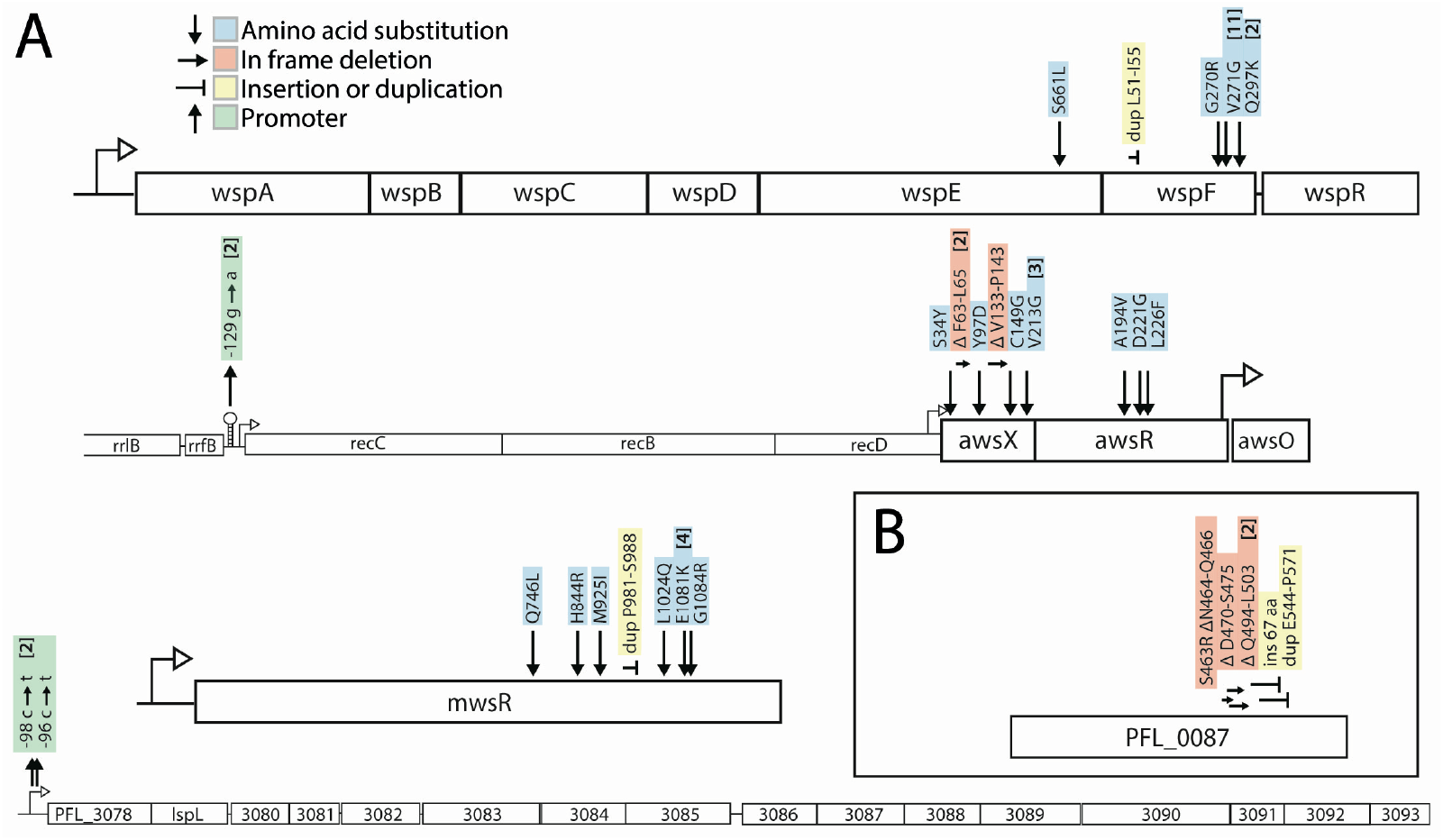
(*A*) Forty-three independent mutants of wild type *Pseudomonas protegens* Pf-5 were isolated after experimental evolution based on their divergent colony morphology and mutations were identified in four operons. Numbers in brackets are the number of independent mutants found. Details are available in S2 Table. (*B*) Experimental evolution with a Δ*wsp* Δ*aws* Δ*mws* triple deletion mutant resulted in WS types with mutations in the DGC PFL_0087/DgcH.

The mathematical null model ((26), S1 Text) successfully predicted that of the three common pathways to WS, Wsp would be the most common (16 mutants) followed by Aws (14) and then Mws (10) (Fig 1E). Mutations were predominately found in the negative regulators WspF (15 mutants) or AwsX (9), but also in interacting proteins WspE (1) and AwsR (3) (Fig 1E). Given that the mutational target size is expected to be smaller for the interacting proteins (Fig 2) this is not surprising. However, no mutations were found in WspA despite a predicted high rate (Fig 1E). The majority (16 out of 22) of the mutated sites or regions was predicted for WspF, WspE, AwsX, AwsR and MwsR (Fig 2), but in most cases they were not identical to those in SBW25 (S2 Table). A further analysis of the likely functional effects of mutations and how they change reaction rates in the model of the genotype-to-phenotype map is available in the S1 Text.

A mutational hot spot was apparent in WspF with 11 out of 15 mutations being identical V271G missense mutations. The previously described mutational hot spots in SBW25 in the *awsX, awsR* and *mwsR* genes (26) appeared absent, demonstrating how mutation rate differences can skew evolutionary outcomes even for closely related species with similar genotype-to-phenotype maps. Mutations in WspA were predicted to be one of the major mutational routes to WS based on the mathematical model (Fig 1E, (26)), but no mutations were found either in this study or in SBW25 (23). However, when the mutational spectrum of WS mutants was determined in the absence of selection for growth at the air-liquid interface, WspA mutants occurred at rates similar to those of WspE and WspF, as predicted by the model, and their low frequency after experimental evolution could be explained by their lower fitness (26). To test if a WspA mutation could cause a WS phenotype in *P. protegens* and measure its fitness, a common deletion mutation found in SBW25 (WspA T293-E299) was introduced into the native gene in Pf-5, resulting in a typical wrinkly colony morphology (Fig 4A).

**Fig. 4.**
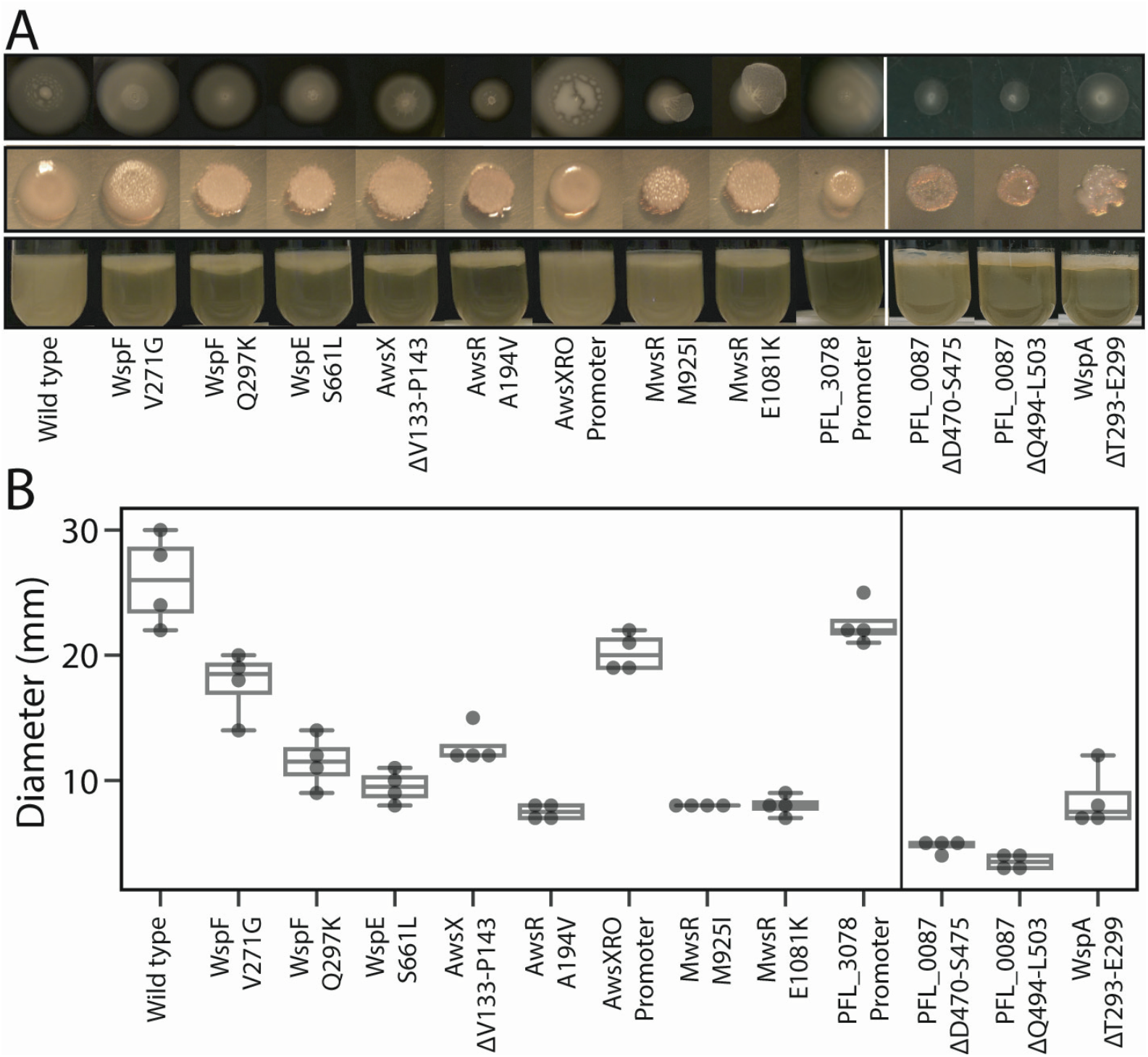
Phenotypic characterization of reconstructed mutants. (*A*) Motility, colony morphology and air-liquid interface colonization of reconstructed representative mutations. (*B*) As expected, motility was significantly reduced for all mutants if the c-di-GMP network was activated, but was only slightly reduced for the AwsXRO promoter capture. Additionally, motility of the PFL_3078 promoter mutant was not significantly reduced, but this was not expected to have increased c-di-GMP levels (one-way ANOVA, F_(12,39)_ = 63.6, p < 0.0001, pairwise differences assessed with student *t*-tests with α = 0.05). Four replicates on two separate plates per strain were used.

To investigate if there are also rare pathways to the WS phenotype, the entire *wsp, aws*, and *mws* operons were deleted and experimental evolution was repeated as previously done for SBW25 (21). Despite inoculating 60 wells with Pf-5 Δ*wsp* Δ*aws* Δ*mws*, as was done with the wild type, WS types were only detected in 7 wells, which could indicate a lower mutation rate to WS when the common pathways are absent. Mutations in the DGC PFL_0087/DgcH accounted for six out of the seven WS types found (Fig 3B). This was also the dominant pathway in the SBW25 Δ*wsp* Δ*aws* Δ*mws* strain where mutations in the corresponding region of PFLU0085 were responsible for 47% of WS mutants.

#### Adaptive mutants colonize the air-liquid interface and most have WS morphology and decreased motility

We evaluated our phenotypic predictions in terms of ability to colonize the air-liquid interface (Fig 1A), colony morphology caused by overproduction of EPS (Fig 1B), and reduced motility (Fig 1C). Due to the difficulties of constructing large numbers of mutants in non-model species, it was necessary here to focus on a select set of diverse mutants. Representative mutations were reconstructed in the wild type using an allelic exchange protocol to determine that the identified mutations were the sole cause of air-liquid interface colonization and colony morphologies, and to exclude the influence of secondary mutations (Fig 4A) before further characterization. Two different phenotypes were observed among the Pf-5 mutants. First, mutants similar to the original WS type in SBW25 had a clear motility defect and mutations in the Wsp, Aws, Mws, and PFL_0087/DgcH pathways (Fig 4A, Fig 4B; one-way ANOVA, F_(12,39)_ = 63.6, p < 0.0001, pairwise differences assessed with student *t*-tests with α = 0.05). The second phenotype observed was less wrinkly and had similar motility as the wild-type (Fig 4A, Fig 4B; one-way ANOVA, F_(12,39)_ = 63.6, p < 0.0001, pairwise differences assessed with student *t*-tests with α = 0.05) and mutations upstream of AwsXRO and the PFL_3078-3093 operon. The phenotypic differences of the AwsXRO putative promoter capture could be due to a lower activation of the c-di-GMP network or by its overexpression being linked to a ribosomal RNA promoter that is highly expressed only under fast growth. That is, as growth rate decreases, the cells are expected to reduce EPS synthesis and increase motility. Mutations in the promoter region of the PFL_3078-3093 operon are not expected to increase c-di-GMP levels, and thus these mutants are not expected to exhibit reduced motility unless expression of the Pap polysaccharide (50) in itself hinders movement.

#### Mutants vary in competitive fitness and rapidly invade wild type populations

In order to evaluate several of our predictions we need to measure the fitness of the mutant under similar conditions as during experimental evolution. This include predictions that the adaptive mutants can invade a wild type population and that the rarity of certain predicted mutants, for example WspA, are due to lower fitness compared to other WS types. Fitness measurements can also provide important data on if the relative fitness effects of mutations are conserved between species. Two types of fitness assays were performed as previously described (21) to measure differences in fitness. The first assay measures “invasion fitness” where the mutant is allowed to invade a wild type population from an initial frequency of 1%. This confirms that the mutations are adaptive and that mutants can colonize the air-liquid interface. The invasion assays showed that all reconstructed mutants could rapidly invade an ancestral wild type population (Fig 5A, S3 Table). Although there were significant differences between selection coefficients of the mutants (one-way ANOVA, F_(12,68)_ = 73.7, p < 0.0001, pairwise differences assessed with student *t*-tests with α = 0.05), no mutant was significantly different from the most common mutant (WspF V271G, two-tailed *t*-test p > 0.01).

**Fig. 5.**
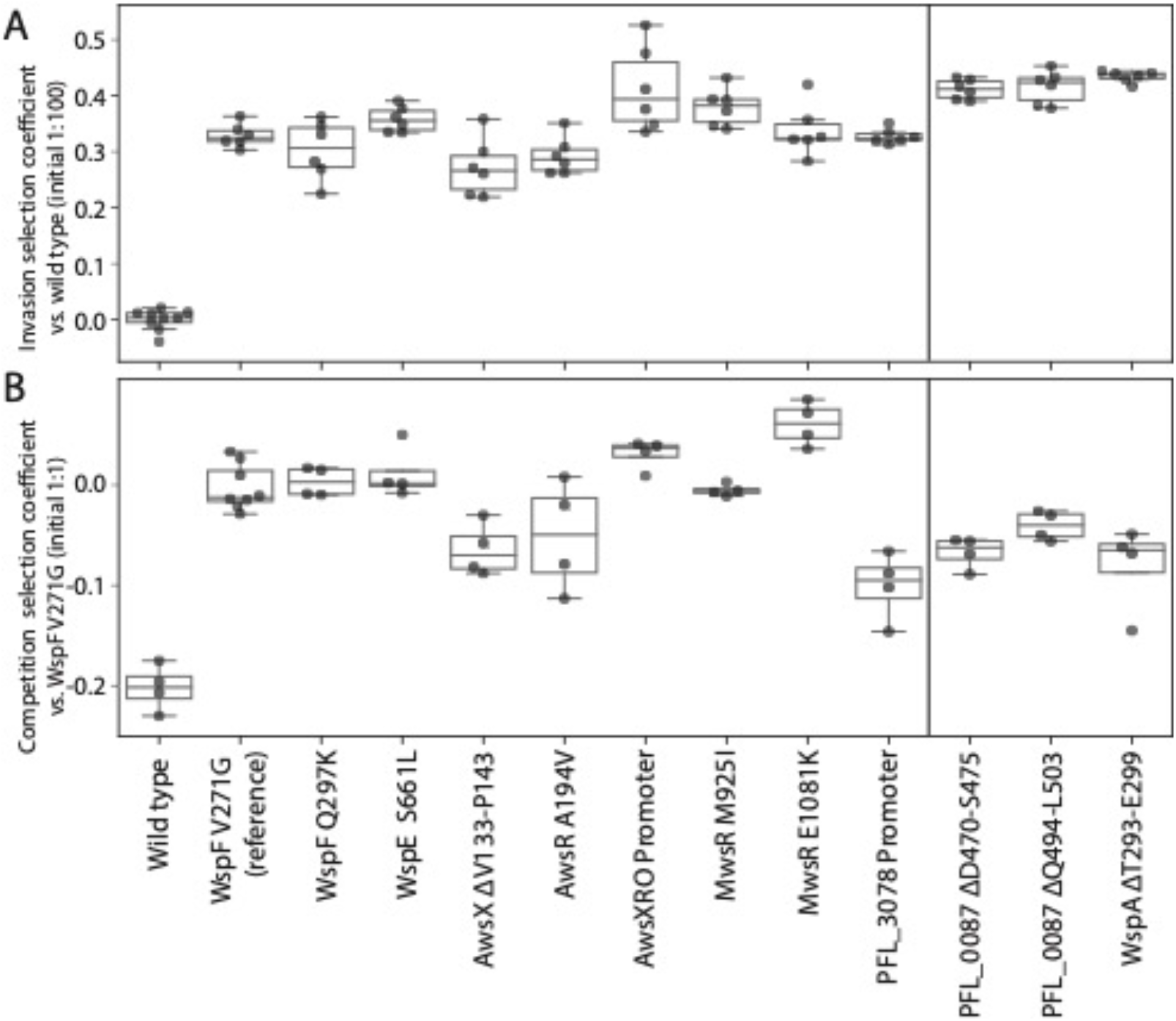
Fitness of reconstructed *P. protegens* Pf-5 WS mutants was measured in pairwise competitions. (*A*) Invasion fitness was measured relative a dominant ancestral wild type strain with a 1:100 initial ratio. All mutations were adaptive and can increase from rare to colonize the air-liquid interface. Six independent competitions were performed for each pair. (*B*) Competition fitness was measured relative the most common WS mutant (WspF V271G) in a 1:1 initial ratio to compare the fitness of different WS mutants and the alternative phenotypic solutions. The mutants with the lowest fitness were the alternative phenotypic solution PFL_3078 promoter mutant and mutants that were not observed after experimental evolution of the wild-type population: PFL_0087 mutants and the constructed WspA mutant (one-way ANOVA, F_(12,43)_ = 25.5, p < 0.0001, pairwise differences assessed with student *t*-tests with α = 0.05). Four independent competitions were performed for each pair.

The second fitness assay measures “competition fitness”. Here, each mutant is mixed 1:1 with the most common WS type (WspF V271G) at the start of the competition (21). The competition assay also showed that the ancestral wild type was rapidly outcompeted by a WS mutant at a 1:1 initial ratio (Fig 5B, S3 Table). There was significant variation in fitness between the WS mutants (one-way ANOVA, F_(12,43)_ = 25.5, p < 0.0001, pairwise differences assessed with student *t*-tests with α = 0.05). The PFL_3078 promoter mutant resulted in the lowest fitness (s = -0.1, two-tailed *t*-test p < 0.0001) meaning that it is expected to be rapidly outcompeted by the WS mutants (Fig 5B). The PFL_0087 mutants that were only found when the common pathways were deleted had lower fitness (two-tailed *t*-tests p = 0.0005, p = 0.001) and this was also true for the constructed WspA mutant (two-tailed *t*-test p = 0.002), which could explain why these were not found in the wild type population after experimental evolution.

#### Pel contributes to AL colonization together with another unknown adhesion factor

WS mutants form stable mats at the air-liquid interface, but the molecular solution to increased cell-cell and cell-surface adhesion needs to be determined to evaluate our phenotypic prediction further. To test if the Pel exopolysaccharide is the main structural component (Fig 1B) used by the different WS mutants of Pf-5, the *pelABCDEFG* operon (PFL_2972-PFL_2978) was deleted from Pf-5 and combined with previously characterized WS mutations and fitness was measured. Both invasion fitness (Fig 6A) and competition fitness (Fig 6B) were significantly lower (two-tailed t-tests p < 0.01) compared to isogenic strains with an intact *pel* operon (Fig 5A, Fig 5B, average fitness of mutants with intact *pel* are plotted as red triangles in Fig 6, S3 Table) except invasion fitness for the AwsX mutant (two-tailed t-tests p = 0.08, one outlier). This suggests that Pel polysaccharide serves as an important structural component for colonizing the air-liquid interface and that Pel production is activated by mutations leading to increased c-di-GMP levels. Although deletion of *pel* operon in WS mutants resulted in less wrinkly colony morphology, it did not result in a smooth ancestral type. Neither did deletion of *pel* abolish the ability to colonize the air liquid interface (Fig 6C) or the ability to invade wild type populations (Fig 6A). This suggests that increased c-di-GMP levels induce production of an additional adhesive component, at least in the absence of *pel*. In SBW25, *pgaABCD* encodes an alternative c-di-GMP-controlled EPS that is used when the cellulose operon is deleted (22). Deletion of the *pgaABCD* operon (PFL_0161-PFL_0164) or the Psl biosynthetic locus (*pslABCDEFGHIJKN*, PFL_4208-PFL4219) in Pf-5 strains with deletion of the *pel* operon combined with WS mutations did not result in a reversion to a wild type phenotype. As expected, if the motility defect observed for WS mutants is primarily caused by high c-di-GMP levels rather than high production of Pel, the motility was also reduced for WS mutants with *pel* deleted (Fig 6D, intact *pel* WS mutants plotted as red triangles).

**Fig. 6.**
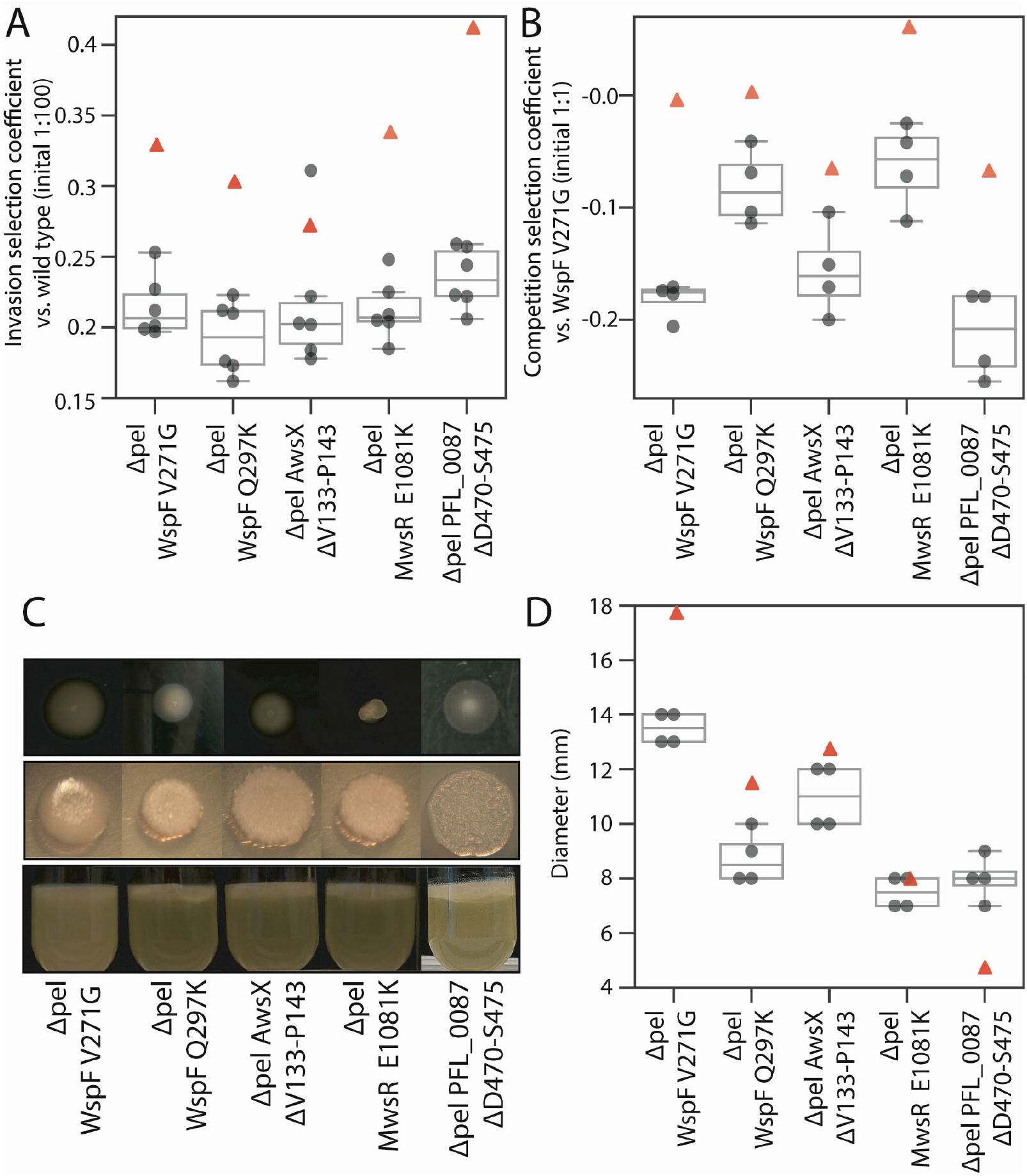
Contribution of *pel* to WS phenotype and fitness. (*A*). Deletion of *pel* in WS mutants reduces invasion fitness (mean values of intact *pel* mutants plotted as red triangle in all plots). Fitness of reconstructed *P. protegens* Pf-5 WS mutants without the *pel* operon was measured in pairwise competitions. Invasion fitness was measured relative a dominant ancestral wild type strain with a 1:100 initial ratio. Six independent competitions were performed for each pair. (*B*) Deletion of *pel* in WS mutants reduces competition fitness. Competition fitness was measured relative the most common WS mutant (WspF V271G) in a 1:1 initial ratio. Four independent competitions were performed for each pair. (*C*) Deletion of *pel* in WS mutants did not result in ancestral smooth colony morphology or loss of ability to colonize the air-liquid interface suggesting a secondary EPS component is produced. (*D*). Deletion of *pel* did not restore motility showing that Pel overproduction is not the cause of the motility defect in WS mutants. Four replicates on two separate plates per strain were used. Deletion of the *pel* operon in the absence of WS mutations did not significantly reduce fitness in invasion against the wild type (0.020 +/-0.050 (SD), two-tailed t-test P = 0.38) or in competition against the wild type (−0.011 +/-0.024 (SD) two-tailed t-test P = 0.49).

### Predictions for other *Pseudomonas* species

Many of our predictions for Pf-5 (Fig 1) are also applicable to other *Pseudomonas* species and in showing that the WS system can be extended to a related species we lay the foundation for a diverse experimental system for testing evolutionary forecasts. This can provide unbiased tests of our ability to predict short-term evolutionary processes and answer fundamental questions about the conservation of genotype-phenotype-fitness maps and mutational biases needed to make general forecasts. Five other *Pseudomonas* species (Fig 7 legend) were chosen to represent the phylogenetic diversity of *Pseudomonas* (S3 Fig) (51) and considering their complement of DGCs and EPSs (Fig 7A, Fig 7B, S4 Table) (50) to make further predictions. All species are obligate aerobes with the exception of the facultative anaerobes *P. aeruginosa* and *P. stutzeri* that are also expected to grow faster with access to oxygen and respond to selection for colonization of the air-liquid interface. These species encode from none to all three of the main DGCs used in SBW25 and only three species contain genes related to cellulose biosynthesis, the main EPS used in SBW25 (Fig 7A, Fig 7B, S4 Table). A summary of predictions at different biological levels is shown in Table 1.

**Table 1.**
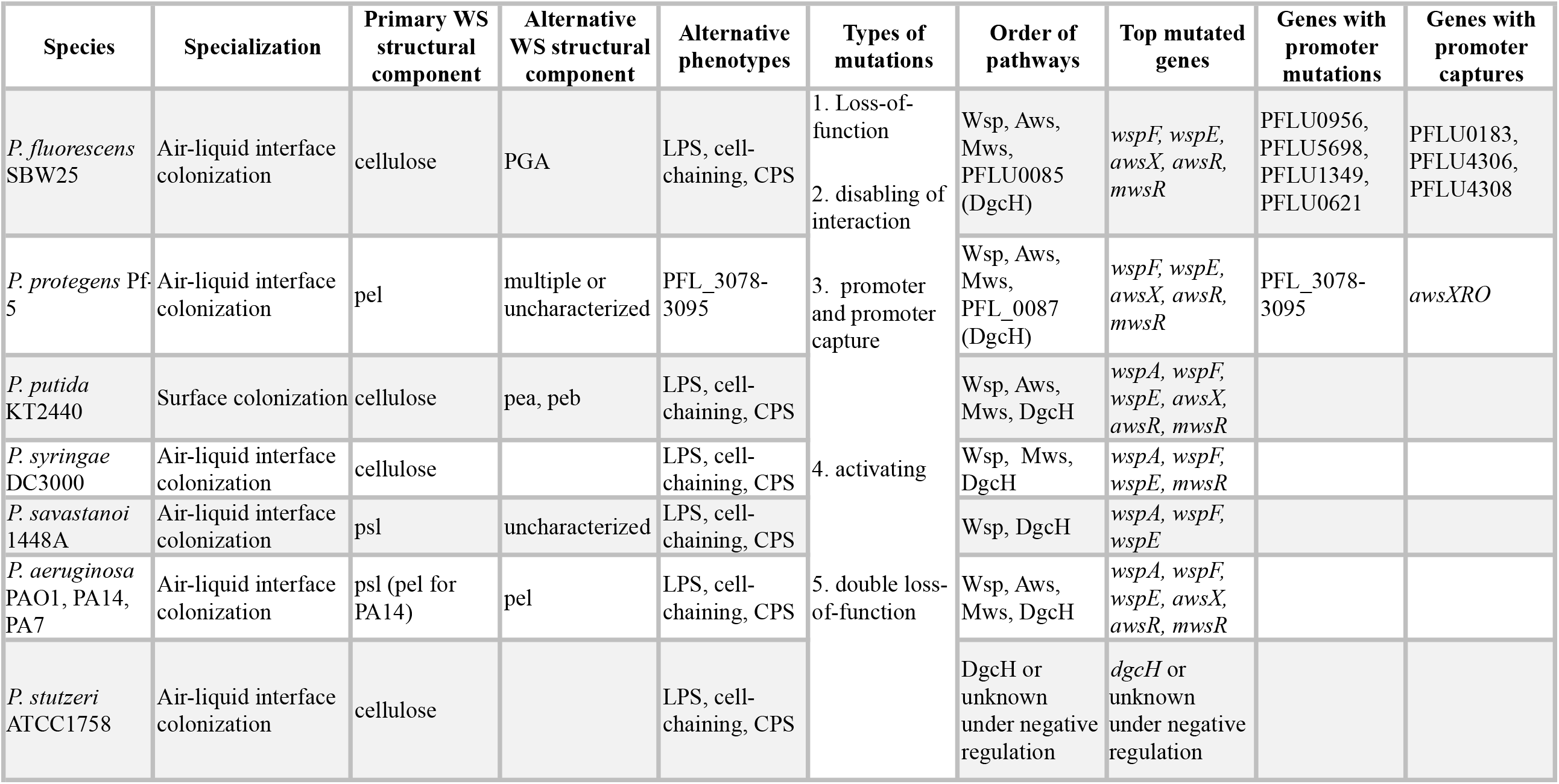
Predictions for other *Pseudomonas* species.

**Fig. 7.**
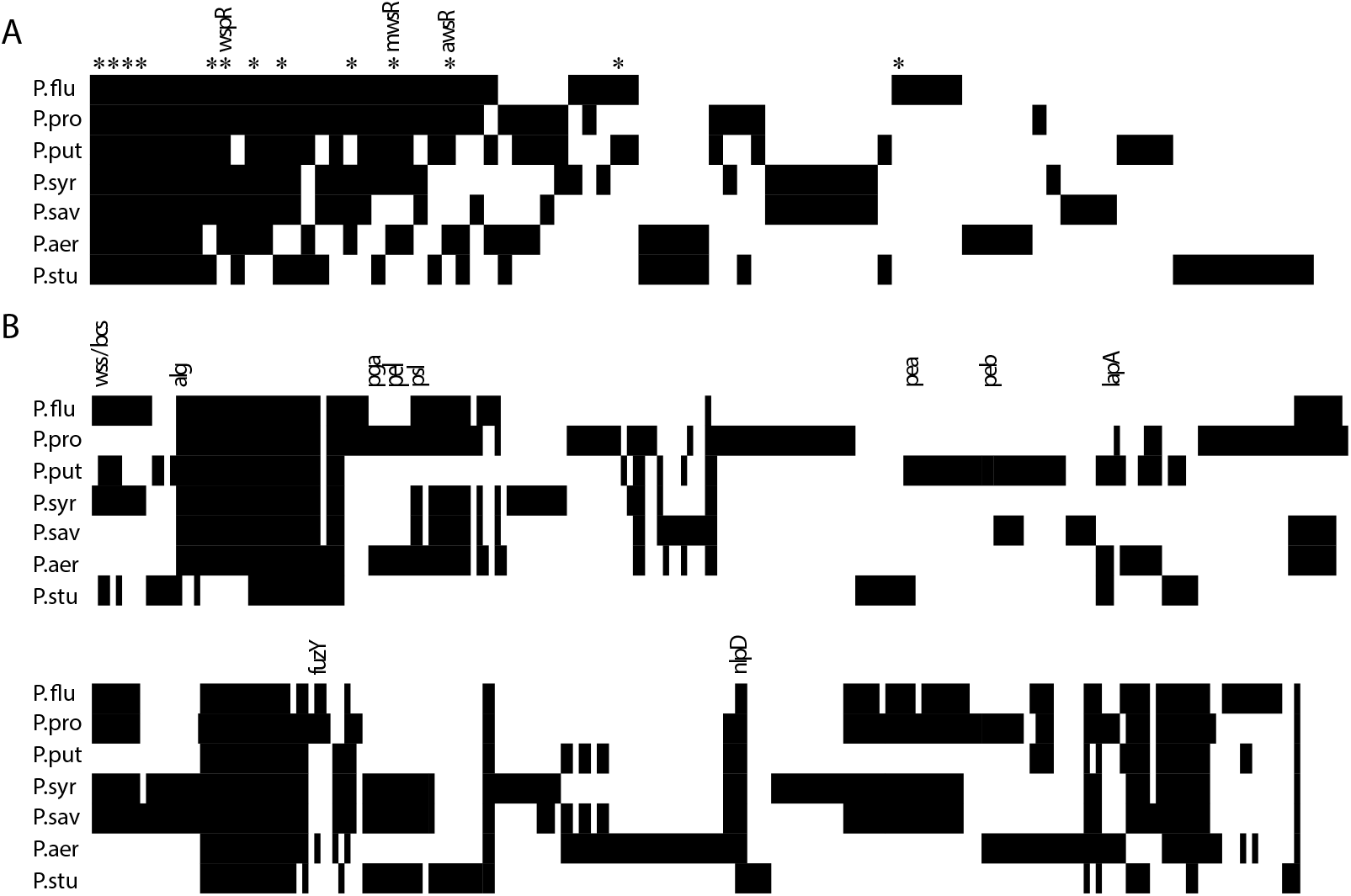
Diversity of DGCs and biofilm-related genes for seven *Pseudomonas* species. (*A*) Five other *Pseudomonas* species (*P. putida* KT2440, *P. syringae* pv. tomato DC3000, *P. savastanoi* pv. phaseolicola 1448A, *P. aeruginosa* PAO1, *P. stutzeri* ATCC 17588) were chosen based on phylogenetic diversity to extend predictions. Including *P. fluorescens* SBW25 and P. *protegens* Pf-5, the seven species encode 251 putative DGCs, divided into 87 different homolog groups of which 8 are present in all genomes. WS mutations in SBW25 have been found affecting 13 of these DGCs (marked with *) with an additional nine that have been detected only in combinations with other mutations. SBW25 and Pf-5 share 33 DGCs with 6 unique for each species. It should be noted that not all DGCs are likely to be catalytically active. (*B*) Diversity of biofilm-related genes including putative EPSs, LPS modification, cell chaining, adhesins and known regulators. The genomes of the *P. aeruginosa* strains PA7, UCBPP-PA14 were also included in the analysis and results were in most cases identical to PAO1 (not shown in Fig 8) with the exception of an absence of homologues for EPS genes *pelA-D* and the DGCs PA2771 in UCBPP-PA14 and PA3343 in PA7. Detailed information is available in S4 Table.

#### Phenotype predictions

Based on the limited data available we predict that cellulose-based biofilms are superior in other species as well and that they will be the primary structural solution when available as for *P. syringae, P. putida* and *P. stutzeri*. For the three species lacking genes for cellulose biosynthesis, we predict that mutants with increased production of other EPSs will have the highest fitness. Based on studies of *P. aeruginosa*, the primary EPS required for pellicle formation at the air-liquid interface in this species is Pel for strain UCBPP-PA14 and Psl for PAO1 and PA7 (52). The genome of *P. savastanoi* lacks genes for biosynthesis of cellulose and Pel and therefore Psl is predicted to be the main structural component.

#### Genetic predictions

The experimental results for Pf-5 largely agrees with predictions and therefore the new data do not give reason for major updates to genetic predictions. The main difference for the other species is that not all commonly used pathways are encoded for all species. *P. savastanoi*, for example, does not have Aws or Mws, so WS mutations are expected to occur at a lower rate and be found mainly in the Wsp pathway. For *P. stutzeri*, none of the three main pathways are available and detailed predictions cannot be made, but it is still possible to predict types of mutations in terms of inactivating mutations followed by promoter mutations and promoter captures. While there is some support of conservation of relative fitness effects of mutations in different genes between Pf-5 and SWB25 in that WspF and WspE mutants are fitter than WspA mutations, despite their different EPS components, it remains to be seen if this is also the case over larger phylogenetic distances. The low frequencies of identical mutations and lack of conservation for mutational hot spots suggest that, at present, a more detailed prediction of exact mutations is not possible and only mutated regions and functional effects can be predicted as done for Pf-5 (Fig 2).

## Discussion

The ability to forecast evolutionary processes could contribute to addressing major societal problems in many key areas including infectious diseases, cancer, climate change and biotechnology (7, 53-55). For many applications, at least in the near future, evolutionary repeatability alone is likely to be sufficient for making forecasts by utilizing large datasets combined with machine learning or statistical models. However, there is a clear advantage of also developing forecasting methods that rely on an understanding of the causes of repeatability because they can be used to test fundamental assumptions about evolutionary processes and provide forecasts for novel situations for which no previous data is available. Such novel situations would also include attempts to control and change evolutionary outcomes by interventions guided by forecasting models. Although there is an extensive theoretical framework for understanding the predictability of evolution, in terms of, for example, fitness landscapes and epistasis (8, 56), predictions about the evolutionary outcome for a specific novel situation are rare and models often use non-accessible parameters or lack explicitly testable hypotheses for failures, such as invoking historical contingency (5).

In this work we explain the basis of our forecasts of experimental evolution, test them experimentally, investigate reasons for failures and extend forecasts to other species to be tested in future studies. We show that this extended wrinkly spreader system has several characteristics that makes it suitable as a model for testing evolutionary forecasts in that the outcomes are not specific to a particular strain or environment, a dominant selective pressure can be established and there are multiple genetic and phenotypic solutions to the adaptive challenge. The selection for increased cell-cell adhesion and biofilm formation is also likely to be relevant ecological traits for natural populations as it can increase antibiotic resistance and immune evasion and decrease protozoan predation (57, 58). However, there are also substantial limitations for this model system in terms of the short time scale of the experiments and the difficulty of incorporating fitness data into models because fitness is frequency-dependent, so there is a clear need for additional model systems where clear predictions can be made and tested.

Our forecasts were accurate in terms of mutants evolving to colonize the air-liquid interface (Fig 1A), use of EPS (Fig 1B), types of mutations (Fig 1D), pathways used (Fig 1E), and mutated regions (Fig 2). These successes are based on the conservation of the fitness of different phenotypes and the conservation of the genotype-to-phenotype map between SBW25 and Pf-5. Determining the reasons for failed forecasts are perhaps more interesting in terms of understanding the limits of our predictive ability and main challenges for moving forward including defining the minimal information needed to make forecasts, what can be predicted *in silico* and what needs to be measured experimentally.

First, strain-specific adaptations pose a major challenge for successful forecasting. Here, the most fit of the two phenotypes used the Pel EPS, which could be predicted from previous data, as well as another yet unidentified structural component. However the other phenotypic solution used a recently identified, though uncharacterized, novel polysaccharide (50) encoded by PFL_3078-3093 for air-liquid interface colonization, demonstrating the difficulty posed by novel strain-specific phenotypic solutions. In contrast to SBW25, where all promoter mutations resulted in up-regulation of DGCs and reduced motility (21), the mutation upstream of PFL3078-3093 demonstrates the possibility of direct transcriptional activation of polysaccharide components that are not under post-translational control of c-di-GMP.

A second challenge is the prediction of relative fitness for different wrinkly spreader mutants. For the multi-protein pathways Wsp and Aws, the mathematical model (S1 Text Model IV in (26)) successfully predicted (Fig 1E) the top mutated proteins except in the case of WspA where no mutations were found. WspA mutants were not found in the original study in SBW25 either (23), but this was shown to be due to lower fitness relative to WspF and WspE mutants rather than a lower mutation rate to WS (26). This is also a likely explanation for the absence of WspA mutants here (Fig 5B), but it remains to be determined if this fitness difference is conserved in other species or if sometimes WspA mutants are more fit. Prediction of the relative fitness effects of adaptive mutants remains a daunting challenge, but it is interesting to note that for both Pf-5 and SBW25 high fitness WS types have mutations in the same proteins (WspF, WspE, MwsR) and low fitness WS mutations were found in WspA, AwsX, AwsR, and PFLU0085/PFL_0087 for both species. If this trend of conservation would extend to other species as well, it could increase the potential of forecasting by partially solving the fundamental problem of how to predict the relative fitness effects of mutations. A possible complication is that relative fitness might be influenced to a great extent by the environment, including the frequency of other adaptive mutants in the population.

The third major challenge is predicting differences in mutation rates as mutational hot spots do not appear to be conserved between SBW25 and Pf-5. Although a mathematical null model that incorporates information about the Wsp, Aws, and Mws molecular networks (Figure 1E, S1 Text, Null model IV in (26)) successfully predicted that Wsp would be the most commonly used pathway to WS (predicted 54%, observed 40%) followed by Aws (predicted 30%, observed 35%), and Mws the most rare (predicted 16%, observed 25%), the high frequency of Wsp mutants seems mainly to be caused by a mutational hot spot in *wspF*. It is also worth noting that direct use of mutation rate data from SBW25 (26) would result in poorer predictions than the mathematical null model due to mutational hot spots in *aws*X, *awsR* and *mwsR* for that species, demonstrating the usefulness of constructing general mechanistic models rather than relying solely on experimental data.

It is possible that an increased mutation rate at promoters (59) is also responsible for the relatively high frequency of mutations upstream of PFL_3078-3093 despite a small mutational target size and the observation that this mutant is rapidly outcompeted WS types (Fig 4B). In SBW25, low fitness phenotypes that colonize the air-liquid interface based on LPS modification or cell-chaining, are observed prior to the rise of WS to high frequencies (22). This is due to the presence of apparent mutational hot spots in these genes, which make these mutants appear early during the growth phase despite their relatively small mutational targets (20, 22, 60). The extent to which mutational hotspots contribute to parallel evolution in experimental evolution studies is unclear. This is partly due to a focus in gene level parallelism and lack of information about exact mutations in databases (61), but the primary problem is the inability to determine the reasons for parallelism in terms of increased mutation rate, differential fitness effects of mutations, and mutational target size (62). In *E. coli*, the most common cases of exact nucleotide parallelism described is associated with IS elements that mediates deletions and duplications though homologous recombination or insertions into gene, while parallelism for point mutations was less common (6, 62). In addition to differences in rates between different base pair substitutions, increased mutation rates associated with nucleotide repeats has also been reported (63, 64). However, general knowledge of such mechanisms do not allow us to *a priori* predict the distribution of mutation rates for a gene or genome and in many cases there is no obvious mechanistic reason for an observed mutational hot spot. It is clear that our current inability to predict mutational hot spots can severely limit predictions across species, even though they can make evolution more deterministic in each single case.

In partially predicting evolutionary outcomes in *P. protegens*, this work lays the foundation for future tests of evolutionary forecasting in related *Pseudomonas* species by clearly stating predictions on several different biological levels - from phenotypes to specific regions of proteins that are likely to be mutated. Currently, given what is already known about the effects of unpredictable mutational biases and differences in fitness between different WS types, many of these forecasts will inevitably fail. However, hopefully they will fail in interesting ways, thereby revealing erroneous assumptions and providing possibilities for iterative improvement of forecasting models. The ability to remove common genetic and phenotypic pathways provides a unique opportunity to also find those pathways that evolution does not commonly use. This is necessary to determine why forecasts fail and update the predictive models for another cycle of prediction, experimental evolution, and mutant characterization that make it possible to use this iterative model to define the information necessary to predict short-term evolutionary processes.

## Materials and methods

### Strains and media

*Pseudomonas protegens* Pf-5 (GenBank: CP000076.1 (65), previously known as *P. fluorescens* Pf-5) and derivatives thereof were used for all experimental evolution and phenotypic characterization. *E. coli* DH5α was used for cloning PCR fragments for genetic engineering. *P. protegens* Pf-5 was grown in tryptic soy broth (Tryptone 17 g, Soytone 3 g, Glucose 2.5 g, NaCl 5 g, K_2_HPO_4_ 2.5 g per liter) supplemented with 10 mM MgSO4 and 0.2% glycerol (TSBGM) for experimental evolution and fitness assays. For comparison, previous studies of *P. fluorescens* SBW25 used King’s medium B for experimental evolution (20 g proteose peptone #3, 1.5 g K_2_HPO_4_, 1.5 g MgSO_4_•7H_2_O and 10 ml glycerol per liter) (21-23, 25, 26, 28). Lysogeny broth (LB) was used during genetic engineering and LB without NaCl and supplemented with 8% sucrose was used for counter-selection of *sacB* marker. Solid media were 1.5% agar added to LB or TSB supplemented with 10 mM MgSO_4_, 0.2% glycerol and 10 mg/l Congo red. Motility assays were conducted in 0.3% agar TSB supplemented with 10 mM MgSO_4_, 0.2% glycerol. Kanamycin was used at 50 mg/l for *E. coli* or 80 mg/l for *P. protegens* and gentamicin at 10 mg/l for *E. coli* or 15 mg/L for *P. protegens*. 100 mg/L nitrofurantoin was used to inhibit growth of *E. coli* donor cells after conjugation. All strains were stored at -80**°**C in LB with 10% DMSO.

### Experimental evolution

Thirty central wells of a deep well plate (polypropylene, 1.1 mL, round walls, Axygen Corning Life Sciences) were inoculated with approximately 10^3^ cells each from independent overnight cultures and incubated at 36**°**C for 5 days without shaking on two different occasions. The wells at the edges of the plate were not used to reduce possible edge effects with increased evaporation and they instead served as contamination controls. Suitable dilutions were plated on TSBGM plates with Congo red after 5 days and incubated at 36**°**C for 48 h. Plates were screened for colonies with a visible difference in colony morphology and one divergent colony per well were randomly selected based only on its position on the agar plate. In total 43 independent mutants were streaked for single cells twice before overnight growth in LB and freezing. An identical protocol was used for the Δ*wsp* Δ*aws* Δ*mws* strain.

### DNA sequencing

Seven mutant strains that did not contain mutations in the *wspF* and *awsX* genes were analyzed by genome resequencing. The strains had mutations in *awsR, mwsR, wspE*, upstream PFL_3078 and in the intergenic region between *rrfB* and *recC* upstream of the awsXRO operon. Genomic DNA was isolated with Genomic DNA Purification Kit (Thermo Fisher). Sequencing libraries were prepared from 1µg DNA using the TruSeq PCRfree DNA sample preparation kit (cat# FC-121-3001/3002, Illumina Inc.) targeting an insert size of 350bp. The library preparation was performed according to the manufacturers’ instructions (guide#15036187). Sequencing was performed with MiSeq (Illumina Inc.) paired-end 300bp read length and v3 sequencing chemistry. Sequencing was performed by the SNP&SEQ Technology Platform in Uppsala. The facility is part of the National Genomics Infrastructure (NGI) Sweden and Science for Life Laboratory. The SNP&SEQ Platform is also supported by the Swedish Research Council and the Knut and Alice Wallenberg Foundation. Sequencing data were analyzed with using Geneious v. 10.2.3 with reads assembled against the *P. protegens* Pf-5 genome sequence (CP000076.1). Sanger sequencing was performed by GATC biotech and used to sequence candidate genes to find adaptive mutations and to confirm reconstructed mutations. Primer sequences are available in S5 Table.

### Reconstruction of mutations

Thirteen mutations representing all candidate genes found as well as PFL_0087 and WspA mutations were reconstructed in the wild type ancestral *P. protegens* Pf-5 to show that they are the cause of the adaptive phenotype and to assay their fitness effects without the risk of secondary mutations that might have occurred during experimental evolution. A two-step allelic replacement protocol was used to transfer the mutation or deletion constructs into the ancestor as previously described (66), but using the mobilizable pK18mobsac suicide plasmid (FJ437239) (full details are available in the S1 Text).

### Fitness assays

Two types of competition fitness assays were performed similarly to previously described assays (21). The first assay measures invasion fitness under 48 h, where a mutant is mixed 1:100 with the wild type ancestor tagged with GFP, mimicking early stages of air-liquid interface colonization where a rare mutant establishes and grows at the surface with no competition from other mutants. The second assay measures competition fitness under 24 h in a 1:1 competition against a reference mutant strain tagged with GFP. We chose the WspF V271G mutant as a reference because it was the most commonly found mutant during experimental evolution. Selection coefficients (s) were calculated as previously described (67) as the change in logarithmic ratio over time according to s = [ln(R(t)/R(0))]/[t], where R is the ratio of mutant to reference and t is the number of generations of the entire population during the experiment. This means that s = 0 is equal fitness, positive is increased fitness, and negative is decreased fitness relative to the reference strain. A full description of the experimental protocol is available in the S1 Text.

### Motility assays

Swimming motility assays were performed in TSBGM plates with 0.3% agar (BD) and the diameter was measured after 24 h of growth at room temperature. Assaying at room temperature, instead at 36 **°**C, reduced in the influence of growth and evaporation to increase reproducibility. Each strain was assayed in duplicates on two different plates, yielding four replicates.

### Bioinformatics analysis of DGCs and EPS genes

Homologs for all DGCs in *P. fluorescens* SBW25 were found using the *Pseudomonas* Ortholog Database (v. 17.2) at Pseudomonas.com (68). Blast-p searches for GGDEF domains were performed to find remaining DGCs in the six *Pseudomonas* species and their homologs again found using the *Pseudomonas* Ortholog Database (69) and manually inspected. Annotations (Pseudomonas.com. DB version 17.2) were also searched for diguanylate cyclase and GGDEF. Not all DGCs found are likely to have diguanylate cyclase activity, but given the difficulties of predicting which of the partly degenerate active sites are likely to be inactive combined with the possibilities of mutational activation during experimental evolution, none were excluded. There is no simple way to find all genes that can function as structural or regulatory genes to allow colonization of the air-liquid interface. Thus, the selection in Fig 7A and S4 table should not be considered complete. Putative EPS genes were found using blast-p searches with sequences of known proteins for exopolysaccharide biosynthesis including cellulose, PGA, Pel, Psl, Pea, Peb, alginate and levan. Homologs were found using the *Pseudomonas* Ortholog Database (69) at Pseudomonas.com (68). Annotations (Pseudomonas.com. DB version 17.2) were also searched for glycosyltransferase, glycosyl transferase, flippase, polysaccharide, lipopolysaccharide, polymerase, biofilm, pili, curli, adhesin and adhesion. Based on previous work in SBW25 and literature searches a few additional genes were added (20-22, 70-73).

## Supporting information

S1 Table Sequence identity

S2 Table Mutations

S3 Table Fitness

S4 Table DGCs and biofilm genes

S5 Table Primers

S6 Table Accession numbers

## Author contributions

Jennifer T. Pentz Investigation, Formal Analysis, Methodology, Visualization, Writing – Original Draft Preparation.

Peter A. Lind, Conceptualization, Funding Acquisition, Investigation, Formal Analysis, Methodology, Resources, Visualization, Supervision, Writing – Original Draft Preparation.

