## Supplementary material for "Forecasting of phenotypic and genetic outcomes of experimental evolution in *Pseudomonas protegens*": S1 text Fig S1-S3

### Supplementary methods

#### Reconstruction of mutations

Thirteen mutations representing all candidate genes found as well as PFL_0087 and WspA mutations were reconstructed in the wild type ancestral *P. protegens* Pf-5 to show that they are the cause of the adaptive phenotype and to be able to assay their fitness effects without the risk of secondary mutations that might have occurred during experimental evolution. A two-step allelic replacement protocol was using to transfer the mutation into the ancestor. First a 1-2 kb fragment surrounding the putative adaptive mutations were amplified using PCR (Phusion High- Fidelity DNA polymerase, Thermo Scientific) and ligated into the multiple cloning site of the mobilizable pK18mobsac suicide plasmid (FJ437239) using standard molecular techniques. The ligation mix was then transformed into competent *E. coli* DH5α using heat shock. After confirmation of correct insert size by PCR the plasmid was transferred to *P. protegens* Pf-5 by conjugation with the donor strain and an *E. coli* strain carrying the conjugation helper plasmid pRK2013. Cultures were grown overnight of the recipient *P. protegens* Pf-5 (20 ml per conjugation at 30°C in LB) and 2 ml each of the donor and helper *E. coli* strains per conjugation at 37°C in LB with kanamycin. The culture of *P. protegens* Pf-5 was heat shocked for 10 minutes at 42°C prior to centrifugation at 4000 rpm for 10 minutes and resuspension in a small volume of LB. Donor and helper cells were collected by centrifugation 4000 rpm for 10 minutes, resuspended in LB, and mixed with the concentrated recipient cells. After another round of centrifugation the conjugation mix was resuspended in 50 µl LB and spread onto several spots on a LB agar plate followed by incubation overnight at 30°C. Each spot of the conjugation mix was scraped from the plate and resuspended in 200 µl LB each and plated on LB agar plates with kanamycin to select for transfer of the plasmid, and nitrofurantoin to prevent growth of the *E. coli* donor and helper cells. The pK18mobsac plasmid has a pBR322 type origin and cannot replicate in *P. protegens* Pf-5. Only cells where the plasmid has integrated into the chromosome by homologous recombination, with the homology provided by the cloned fragment, can grow in the presence of kanamycin. After streaking for single cells on LB agar plates with kanamycin, the *P. protegens* Pf-5 strains with integrated plasmids were grown overnight in LB at 30°C without antibiotics to allow for double crossover homologous recombination resulting in loss of the integrated plasmid. The plasmid also contains the *sacB* marker conferring sucrose sensitivity, which allows for counter-selection by plating on LB agar plates with sucrose. Sucrose resistant colonies were checked for loss of the kanamycin marker and DNA sequencing of the cloned region to find strains with the reconstructed mutation and no other mutations.

Deletion of the *wsp*, *aws*, *mws*, *pelABCDEFG* (PFL_2972-PFL_2978), *pgaABCD* (PFL_0161-PFL_0164) and *pslABCDEFGHIJKN* (PFL_4208-PFL4219) regions was accomplished using the same two-step allelic exchange protocol using SOE-PCR to generate a fragment surrounding the operon as previously described (Ferguson, et al. 2013; Farr 2015; Lind, et al. 2017). Gene synthesis (Thermo Fisher) was used to make DNA fragments used for deletion of PFL_0161-PFL_0164 and WspA T293-E299. Primer sequences are available in supplementary table S5.

**Fitness assays**

Two types of competition fitness assays were performed similarly to previously described assays (Lind, et al. 2015). The first assay measures invasion fitness, where a mutant is mixed 1:100 with the wild type ancestor, simulating early stages of air-liquid interface colonization where a rare mutant establishes and grows at the surface with no competition from other mutants. The second assay measures competition fitness in a 1:1 competition against a reference mutant strain. We chose the WspF V271G mutant because it was the most commonly found mutant during experimental evolution and thus is highly successful, either because of a high rate of emergence, *e.g.*, a mutational hot spot, or higher fitness than most other WS mutants. In addition, the WspF V271G mutant has a temperature sensitive colony morphology phenotype that it is highly wrinkly at 30°C, but has a very mild phenotype when grown at room temperature, allowing it to be distinguishable from both the smooth ancestor and all other wrinkly mutants isolated here.

Fluorescent reference strains of the wild type ancestor and the WspF V271G mutants were created using a miniTn7 transposon (miniTn7(Gm) PA1/04/03 Gfp.AAV-a) (Lambertsen, et al. 2004) that allows integration at a defined locus (attTn7) in the chromosome. This allows the colonies to be distinguished not only by morphology, but by fluorescing under blue/UV light, and confers resistance to gentamicin. This provides a way to ascertain that secondary adaptive mutants that might occur during the competition experiment do not bias the results (for example the ancestor could evolve WS types or a WS mutant can evolve to cheat on the other type by inactivation of EPS production or reduced c-di-GMP signalling). Introduction of the transposon into *P. protegens* Pf-5 was performed by tri-parental conjugation from *E. coli* with helper plasmids pRK2013 (conjugation helper) and pUX-BF13 (containing the transposase genes) using the same conjugation protocol described above.

The invasion assay was performed by mixing shaken overnight cultures of the competitor 1:100 with the GFP-labeled reference ancestor followed by 1000-fold dilution and static incubation at 36°C for 48 h in TSBGM medium in deep well plates (1 ml per well, using only the central 60 wells). For the competition assay, the GFP-labeled reference strain WspF 271G was mixed 1:1 with the competitor and diluted 6-fold and grown for 4 h (shaken at 30°C), before plating to determine initial ratios, to ensure the cells were in a similar physiological state at the start of the competition. The competition cultures were then diluted 1000-fold in TSBGM medium and grown in deep well plates (1 ml per well, using only the central 60 wells) static for 24 h at 36°C. Selection coefficients (s) were calculated as previously described (Dykhuizen 1990), where s = 0 is equal fitness, positive is increased fitness, and negative is decreased fitness relative to the reference strain. Briefly s is calculated as the change in logarithmic ratio over time according to $s = [\ln(R(t)/R(0))]/[t]$, where R is the ratio of mutant to reference and t is the number of generations of the entire population during the experiment (estimated from viable counts). The cost of the fluorescent marker were calculated from control competitions where the GFP-labeled reference strains (wild type and WspF V271G) were competed against isogenic strains without the marker and included in each plate under identical conditions during the fitness assays and used to adjust the selection coefficients to compensate for the cost. The competition and invasion fitness assays were designed to measure two different aspects of fitness likely to be relevant in the experimental evolution experiment based on previous work (Lind, et al. 2015, 2017; Lind, et al. 2019). The competition assays measures the mutants ability to compete at the air-liquid interface with a highly successful mutant and the invasion assay measures the ability to colonise the air-liquid interface in the near absence of other adaptive mutants. Both these assays are conducted over shorter time scales (24 h and 48 h) compared to the experimental evolution experiment (5 days) to reduce the impact of secondary mutants arising during the experiment, which makes it impossible to assay mutants with low fitness. The final population sizes are similar in these assays and the experimental evolution experiment, because the assays are started with a 1000-fold higher inoculum, which is important to reduce the effects of secondary mutants during the experiment.

Four replicates were used to measure competition fitness and six replicates for the invasion assay based on previous experience. A single colony of each competition strain was inoculated for overnight culture for each replicate and mixed with a reference strain. No outliers were excluded. If a large fraction of the colonies (>5%) display a phenotype caused a secondary mutation during the fitness assay these would be excluded, for example if the wild type strain marked with GFP display a WS phenotype for the invasion assay. However, this did not occur for any of the replicates.

#### Modelling of Wsp, Aws and Mws pathways

The molecular networks of the three main pathways (Wsp, Aws and Mws) were modeled as a system of ordinary differential equations and this work is fully described as Model IV in (Lind, et al. 2019) and a general description of the method is available in (Libby and Lind 2019). The model allows us to compute the relative likelihood that a mutation will translate in to phenotypic change for the three main pathways to WS. Here we only use predictions from model IV and assume that mutations that produce disabling changes that reduce reaction rates are ten times more common than enabling changes. We also assume that the genotype-to-phenotype map is conserved and the modeled functions of the proteins in the Wsp, Aws and Mws are conserved, which means that there are no changes to the system of differential equations.  This does not necessarily mean that all aspects of the molecular details of the systems are conserved and they can also, for example, respond to different environmental signals. Importantly, the model does not take into account the nature of the phenotype produced, in terms of exopolysaccharide used, as it computes the change to an indicator of phenotypic change, which is the active form of the DGC for each network. Therefore results from the model can be directly used for prediction in Pf-5.

The first step is to describe the biochemical reactions of the molecular network by a set of ordinary differential equations (Fig S1) that describe the reactions kinetics of the indicator of phenotypic change, in this case the active form of the DGC producing the c-di-GMP signal (R* for Wsp, RR for Aws and D* for Mws).

### A Wsp

1. Methylated WspC activates WspABD by methylation
2. Phosphorylated WspF inactivates methylated WspABD by demethylation
3. Signal S activate s methylated WspABD by autophosphorylation
4. Methylated and phosphorylated WspABD activate s WspE by phosphorylation
5. Phosphorylated WspE phosphorylate s WspR to an active form (indicator)
6. Phosphorylated WspE phosphorylate s WspF
7. WspR is degraded or bound competitor at constant concentration

1. $C\# + ABD \xrightarrow{r_1} ABD\#$
2. $F^* + ABD\# \xrightarrow{r_2} F + ABD$
3. $ABD + S \xrightarrow{r_3} ABD\#^*$
4. $ABD\#^* + E \xrightarrow{r_4} ABD\# + E^*$
5. $E^* + R \xrightarrow{r_5} R^* + E$
6. $E^* + F \xrightarrow{r_6} F^* + E$
7. $R \xrightarrow{.01} \varnothing$

$$\frac{\partial[ABD]}{\partial t} = r_2[F^*][ABD\#] - r_1[ABD][C\#]$$
$$\frac{\partial[ABD\#]}{\partial t} = r_1[ABD][C\#] - r_2[F^*][ABD\#] - r_3S[ABD\#] + r_4[E][ABD\#^*]$$
$$\frac{\partial[ABD\#^*]}{\partial t} = r_3S[ABD\#] - r_4[E][ABD\#^*]$$
$$\frac{\partial[E^*]}{\partial t} = r_4[E][ABD\#^*] - r_5[E^*][R] - r_6[E^*][F]$$
$$\frac{\partial[R^*]}{\partial t} = r_5[R][E^*]$$
$$\frac{\partial[E]}{dt} = r_6[E^*][F] - r_4[E][ABD\#^*] + r_5[R][E^*]$$
$$\frac{\partial[F]}{\partial t} = r_2[F^*][ABD\#] - r_6[E^*][F]$$
$$\frac{\partial[F^*]}{d\partial t} = r_6[E^*][F] - r_2[F^*][ABD\#]$$
$$\frac{\partial[R]}{\partial t} = -r_5[R][E^*] - .01[R]$$

### B Aws

1. Signal S activates AwsO
2. Activated AwsO binds to AwsX to form a complex that sequesters AwsX
3. AwsX binds AwsR to form a complex that sequesters AwsX
4. AwsR dimerizes to active form (indicator)

1. $S + O \xrightarrow{r_1} O^\wedge$
2. $X + O^\wedge \xrightarrow{r_2} OX$
3. $X + R \xrightarrow{r_3} XR$
4. $R + R \xrightarrow{r_4} RR$

$$\frac{\partial[X]}{\partial t} = -r_2[X][O^\wedge] - r_3[X][R]$$
$$\frac{\partial[XR]}{\partial t} = r_3[X][R]$$
$$\frac{\partial[O]}{\partial t} = -r_1S[O]$$
$$\frac{\partial[O^\wedge]}{\partial t} = r_1S[O] - r_2[X][O^\wedge]$$
$$\frac{\partial[OX]}{\partial t} = r_2[X][O^\wedge]$$
$$\frac{\partial[R]}{\partial t} = -r_3[X][R] - r_4[R][R]$$
$$\frac{\partial[RR]}{\partial t} = r_4[R][R]$$

### C Mws

1. Signal S activates DGC domain (indicator)
2. Activated DGC interacts with PDE to inactivate it
3. Activated DGC interacts with C and to form alternative complex DC

1. $S + D \xrightarrow{r_1} D^*$
2. $D^* + P \xrightarrow{r_2} DP$
3. $D^* + C \xrightarrow{r_3} DC$

$$\frac{\partial[D]}{\partial t} = -r_1[D]S$$
$$\frac{\partial[D^*]}{\partial t} = r_1[D]S - r_2[D^*][P] - r_3[D^*][C]$$
$$\frac{\partial[P]}{\partial t} = -r_2[D^*][P]$$
$$\frac{\partial[DP]}{\partial t} = r_2[D^*][P]$$
$$\frac{\partial[C]}{\partial t} = -r_3[D^*][C]$$
$$\frac{\partial[DC]}{\partial t} = r_3[D^*][C]$$

**Figure S1** – Description of mathematical model of Wsp, Aws and Mws (adapted from (Lind, et al. 2019).

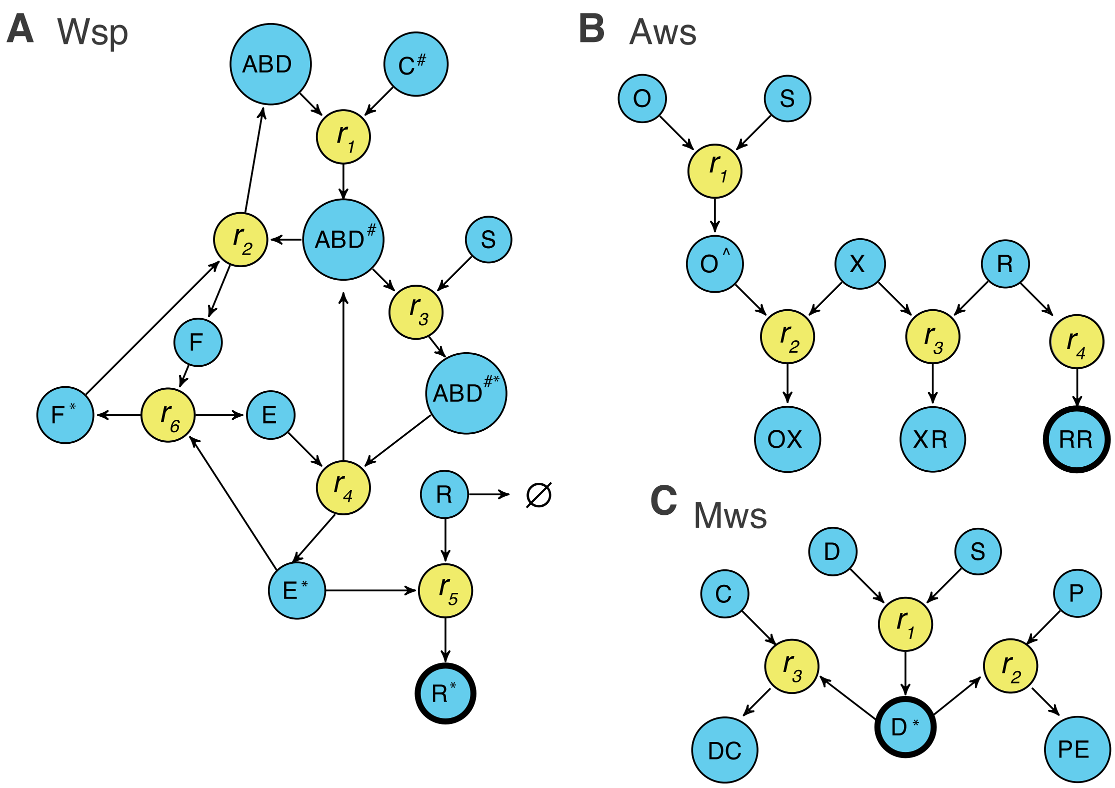


**Figure S2** – Reaction networks of Wsp, Aws and Mws (adapted from (Lind, et al. 2019). The indicator of phenotypic change for each network (R* for Wsp, RR for Aws and D* for Mws) is shown in as bold edged circles.

To solve the system of differential equations knowledge of reaction rates and concentrations are needed. However, it is rarely possible to experimentally assay reaction rates and concentrations for proteins in regulatory networks. This modeling method was designed to produce useful predictions with limited knowledge of only of functional interaction network underlying a phenotype. To do so we choose uniform parameter distributions for initial concentrations U[0,10] and reaction rates $10^{U[-2,2]}$ for the interactions between components. Distributions for effects to reaction rates were chosen as $10^{U[-2,0]}$ for disabling changes and $10^{U[0,2]}$ for enabling changes.

The aim of model is to determine how the concentration of an indicator, in this case the active form of the DGC producing the c-di-GMP signal, change when reaction rates are changed up or down due to mutations. In order to link this to a phenotypic change we set a phenotypic threshold, compared to a previously determined baseline level, that here represent the wrinkly spreader phenotype. Numerical simulations were performed to repeatedly sample from the distributions of reaction rates, initial concentrations and magnitude of effect allows computation of the probability that each set of particular changes to reaction rates produce a wrinkly spreader.

The model also allows computation of the probability that changes to specific reaction rates in a network produce a wrinkly spreader. Changes to a reaction rate can be caused by mutational changes in all participating proteins and here we simply assume that mutations are equally likely in either one. An example of this for the Wsp network is to assume that disabling changes to $r_6$ occur with equal probability by mutations in either WspF or WspE. With knowledge of protein function this can be further modulated to include a predicted target size and an unequal distribution for the proteins in the interaction. However this did not improve predictions in (Lind, et al. 2019), where equal numbers of mutations were found in WspA, WspE and WspF as predicted by the model, but where WspF is expected to have a larger target size.

### Analysis of mutational effects on reaction rates

An important aspect of testing the model is also to determine the effects on protein function of the mutations, which requires an analysis of individual mutations. For example mutations in WspA could produce WS types by either disabling mutations affecting $r_2$ or enabling changes in $r_4$. The molecular effects of the mutations found here are unknown, but knowledge from SBW25 and *P. aeruginosa* and their positions in protein structure allowed limited analyses of likely functional effects. Inactivating mutations in the negative regulator WspF were predicted to be either indels or missense mutations in four specific regions causing disabling changes to $r_2$ and $r_6$ in the model. Mutations were found in two of the predicted regions, one in the vicinity to the methylesterase active site where mutations are predicted to cause disrupt the catalytic site and reduce $r_2$ and the other one directly disrupting the phosphorylation active site in the signal receiver domain thereby reducing $r_6$. No mutations were found in the surface exposed regions hypothesized to be involved in interactions with WspA and WspE, which could be due to differences in function between SBW25 and Pf-5 or simply that they appear at lower frequency and would be detected if additional mutations were isolated. The sole mutation in WspE is, as predicted, located in the direct vicinity of the phosphorylation active site and is predicted to reduce $r_6$. Mutations in AwsX were amino acid substitutions throughout the gene as well as in frame deletions inactivating the gene as predicted that is expected to produce their phenotypic effect by reducing $r_3$, although $r_3$ would also be reduced if function is completely disabled. Mutations in AwsR and MwsR were also found in predicted regions, but no mutations were found in the small periplasmic region of AwsR, which is the most commonly targeted region in SBW25 where it is likely to reduce $r_3$. Known mutational hot spots in *awsX*, *awsR* and *mwsR* in SBW25 (Lind, et al. 2019) were not conserved in Pf-5 resulting in divergent spectra of mutations, while mutated regions and predicted functional effects remain conserved between the two species. Little is known about the molecular function of the putative DGC encoded by PFL_0087/PFLU0085, but it is clear that a multitude of amino acid substitutions, deletions and insertions in a more than 40 amino acids long region can lead to WS (Lind, et al. 2015). Thus it functions as a small intragenic negative regulator region that have been proposed to be involved in oligomerization (Lind 2019) and loss of this interaction results in constitutive activation of c-di-GMP production.

### Construction of Multi-Locus Sequence Analysis phylogenetic tree

The sequences of the four housekeeping genes 16sRNA, *gyrB*, *rpoB*, and *rpoD* were retrieved for 15 popular *Pseudomonas* species using available databases (GenBank accession numbers are listed in Supplementary Table S6). The sequences were subsequently concatenated in the following order: 16sRNA (1543 nt), *gyrB* (2430 nt), *rpoB* (4096 nt), and *rpoD* (1882 nt) (Mulet, et al. 2010) and a multi-locus alignment was created using the MEGA X software (version 10.1.7) (Kumar, et al. 2018) by implementing the MUSCLE algorithm. A phylogenetic tree was created using the maximum likelihood method and the Tamura-Nei model of nucleotide substitution (Mulet, et al. 2010).

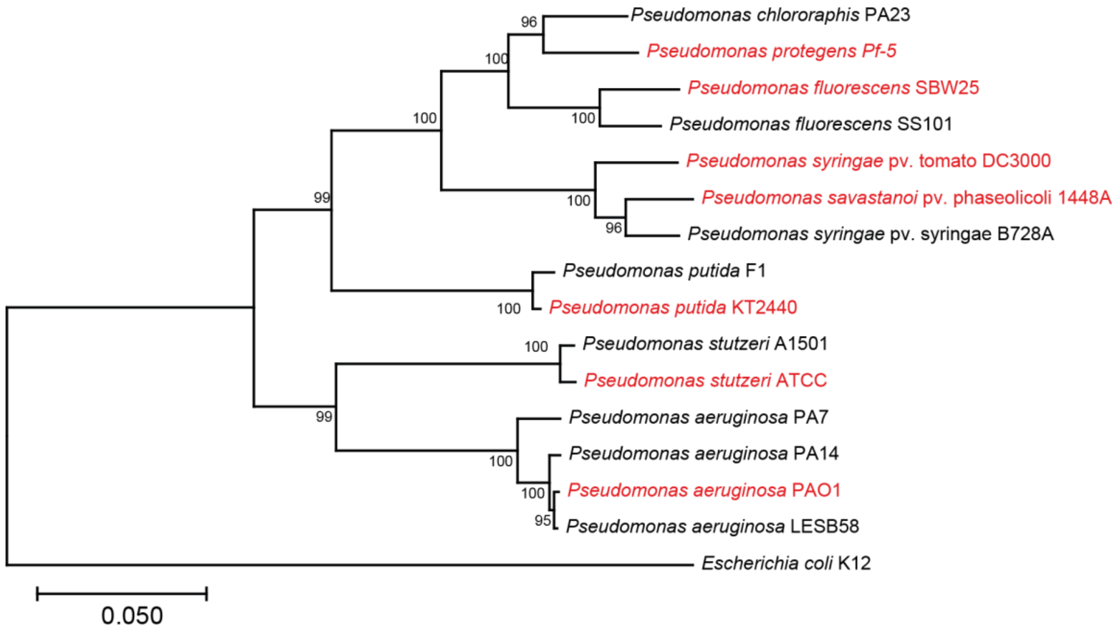


**Figure S3** – Phylogeny of core genes for *Pseudomonas* species selected for evolutionary forecasting. Phylogenetic tree of 15 commonly studied *Pseudomonas* species based on the concatenated sequences of four genes (16sRNA, *gyrB*, *rpoB*, *rpoD*). Numbers at branch nodes indicate bootstrap values of 1000 replicates. The seven species used for further analysis of DGC and biofilm-related genetic diversity (Fig 7A, 7B) are labeled in red ((*P. fluorescens* SBW25, P. *protegens* Pf-5, *P. putida* KT2440, *P. syringae* pv. tomato DC3000, *P. savastanoi* pv. phaseolicola 1448A, *P. aeruginosa* PAO1, *P. stutzeri* ATCC 17588).
